# Copyright and the Use of Images as Biodiversity Data

**DOI:** 10.1101/087015

**Authors:** Willi Egloff, Donat Agosti, Puneet Kishor, David Patterson, Jeremy A. Miller

## Abstract

Taxonomy is the discipline responsible for charting the world’s organismic diversity, understanding ancestor/descendant relationships, and organizing all species according to a unified taxonomic classification system. Taxonomists document the attributes (characters) of organisms, with emphasis on those can be used to distinguish species from each other. Character information is compiled in the scientific literature as text, tables, and images. The information is presented according to conventions that vary among taxonomic domains; such conventions facilitate comparison among similar species, even when descriptions are published by different authors.

There is considerable uncertainty within the taxonomic community as to how to re-use images that were included in taxonomic publications, especially in regard to whether copyright applies. This article deals with the principles and application of copyright law, database protection, and protection against unfair competition, as applied to images. We conclude that copyright does not apply to most images in taxonomic literature because they are presented in a standardized way and lack the creativity that is required to qualify as 'copyrightable works'. There are exceptions, such as wildlife photographs, drawings and artwork produced in a distinctive individual form and intended for other than comparative purposes (such as visual art). Further exceptions may apply to collections of images that qualify as a database in the sense of European database protection law. In a few European countries, there is legal protection for photographs that do not qualify as works in the usual sense of copyright. It follows that most images found in taxonomic literature can be re-used for research or many other purposes without seeking permission, regardless of any copyright declaration. In observance of ethical and scholarly standards, re-users are expected to cite the author and original source of any image that they use.

## 2. Introduction

Communication is a key part of science. Through access to prior scientific results and through communication of new results, we collectively assemble a better understanding of the world than can be achieved by individuals working in isolation. Communication allows sceptics to assess prior work, repeating the work when warranted. Scientific communication is most reliably achieved by the publication of articles in peer-reviewed journals. It is widely accepted that peer-review helps to ensure that each publication meets community standards of integrity, novelty, conforms to general scientific principles and to the standards and best practices of the relevant scientific domain [1-3].

In order to build on prior results, science is best presented in a standardized way. Publications begin with general background that provides context and identifies the most relevant prior work. Methods of experimental setup and data collection are reported in a dedicated block of text that may be referred to as ‘Materials and Methods’ or a similar heading. New information is presented in the “Results’ section, and their significance is discussed in the context of prior work and current understanding in the ‘Discussion’ section. In the ‘Results’, most measurements are given in internationally standardized units, and may be represented in charts and diagrams.

The advent of the Internet has been followed by the emergence of standard formats for digital data. Data standards include the <u>FASTA</u> format for protein and DNA sequence data, IUPAC/IUB codes for referring to amino acids and nucleotides [4], and <u>Darwin Core</u> for occurrence records [5]. These standards are key to the large-scale synthesis of biodiversity knowledge that has been referred to as a knowledge graph [6].

The spectrum of biodiversity that manifests in the form of different species is the subject matter of taxonomy. Since the first accepted contributions to taxonomy [7-9], taxonomic publications have contained taxonomic treatments. Treatments address the identity of a taxon using a scientific name within a hierarchical classification, list characteristics that define the taxon and distinguish it from all others, report where the taxon has been found, and cite earlier publications with content on that taxon [10]. Community standards as to how this information is expressed, enforced in part by peer review, make it possible for multiple independent researchers to work collaboratively to assemble a unified understanding of life [11].

Contributions to taxonomy may take the form of a taxonomic revision, containing treatments of all species in a supraspecific taxonomic group such as a genus or subfamily. Publications may be geographically limited (to a country, region, continent) or be global in scope. Publications may describe one or a small number of new taxa, or add and refine knowledge regarding a taxon that was described previously. Over time, all taxonomic groups receive contributions from multiple researchers working independently. The conventions of scholarship demand that all relevant previous work be cited. Although this is rarely the case in science, taxonomists are especially diligent in this regard and, ideally, are attentive to ALL previous treatments of a taxon [11]. Elsewhere, we [12] have presented the case that much of the text in taxonomic treatments is not eligible for copyright protection, introducing the ‘Blue List’ to summarize classes of relevant information.

Taxonomic treatments must be published on paper or in electronic form [13, 14]. As to quantity, we do not know how many taxonomic treatments have been published in books and journals as the domain is not sharply defined, with taxonomy grading into ecology, geology, geography, molecular processes, cosmology, and other disciplines. Thomson Reuters specialize in indexing articles about Biology and (at the time of writing) Biosis Previews covers more than 5,200 journals (over 21 million records), Biological Abstracts indexes over 4,200 journals (more than 12 million records), and Zoological Record indexes more than 5000 journals with 3.5 million records (wokinfo.com/media/pdf/BIOSIS_FS.pdf). The Biodiversity Heritage Library has digitized (at the time of writing) and indexed almost 200,000 volumes (more than 50 million pages, perhaps a tenth of all of pages relevant to biodiversity). Only a fraction of these items relate to taxonomy.

## 3. Images as a form of biodiversity data

The identification and diagnostic aspects of taxonomy require researchers to focus on attributes (also known as features, characters, or character states) that differ in some way between taxa. The accounts of those attributes are achieved through a combination of text and images (and increasingly other kinds of content). The presentation is explicitly intended to allow comparison with similar organisms, facilitating the task of pointing to or comparing distinguishing features. To achieve this, images typically depict an organism (in whole or selected parts) with a particular orientation and rendered in a particular style to highlight certain details.

### 3.1 Achieving standard approaches

With multiple independent researchers contributing knowledge to a taxonomic group, communities tend to adopt the same views and formats to better communicate with each other. Scientific illustrators are taught to be aware of conventions operating within the scientific discipline to which they are contributing. “Maintaining consistent conventions permits the work of several illustrators to be easily compared and ensures that an illustration will be 'read' properly” [15]. In digital imaging, parameters such as lightning, optical, and specimen orientation are kept consistent. Distributed collaborative projects such as <u>AntWeb</u> have explicit standards and instructions for creating digital images of standard views [16]. When executed according to the protocol, images and data contributed to the site will be comparable regardless of the supplier (see antweb.org/documentation.do). Standard formats are used to facilitate the transfer and sharing of data [5, 17]. Standards in scientific imaging minimize creative variation to ensure that the subject is represented in a consistent way and can be integrated into the corpus of scientific literature. Because of the need to comply with standards, we argue that such images lack “sufficient creativity”, the central criterion used to determine if an illustration qualifies as a “work” in the sense of copyright law.

The combination of structured text and standard view images in taxonomic treatments is a mechanism for documenting facts [11]. The approach is not expressive in the sense that creative writing and visual arts are. This contrasts with other representations of natural history, such as wildlife illustrations created as pieces of commercial art [15], and also with examples of expressive creativity that occasionally appear in scientific literature (Fig. 1). Such works are excluded from the rights arguments made here.

**Fig. 1.**
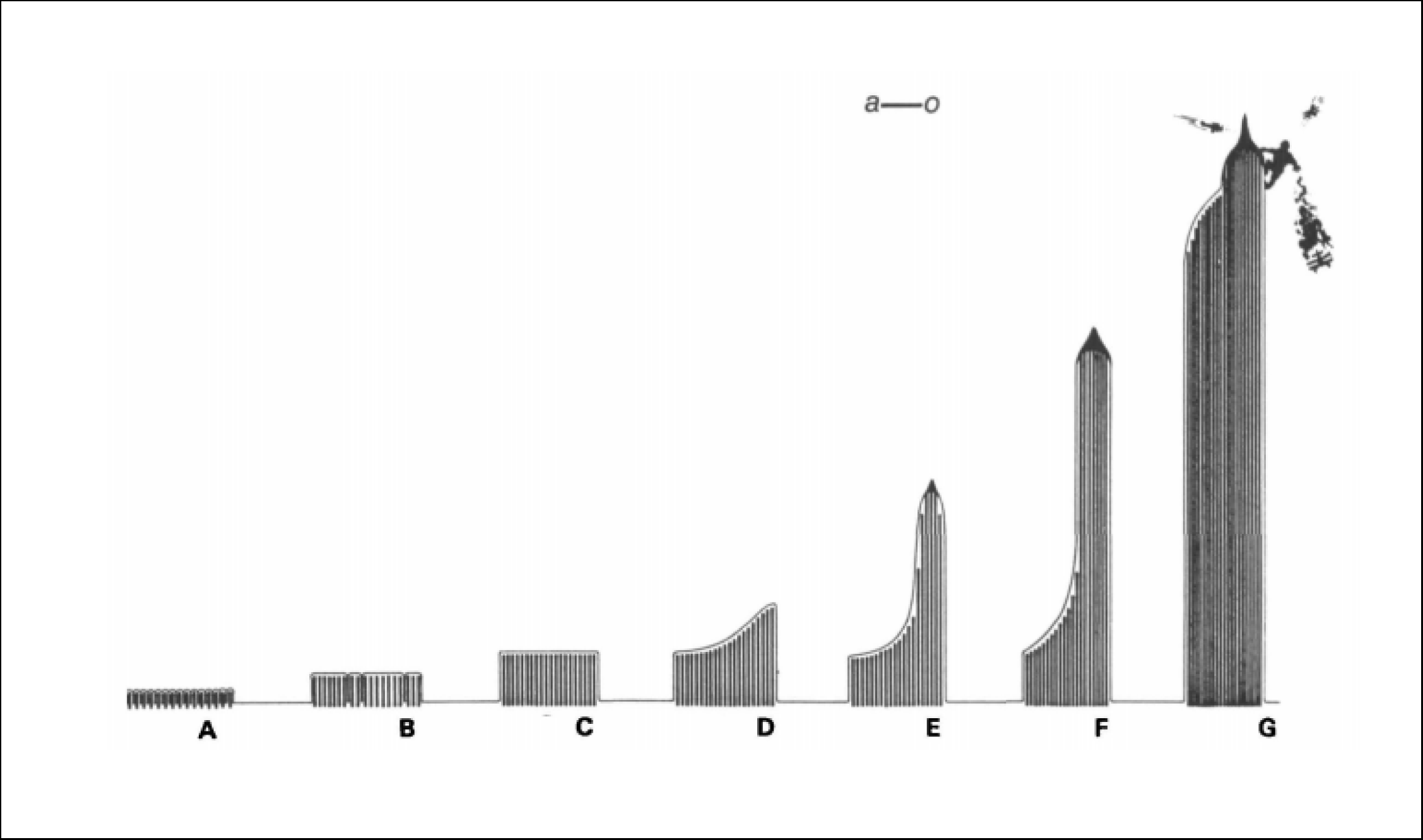
Series of diagrams showing the development of subcellular organelles in a ctenophore. In a touch of creative whimsy, the authors have added King Kong battling aircraft atop the fully developed organelle, which resembles a skyscraper. From Tamm and Tamm 1988 [18].

### 3.2 Consistency over time

Taxonomy began as a scientific discipline in the middle of the 18th century. Botany and zoology designate different works of Carl Linnaeus as the starting points for their respective taxonomic domains: Species Plantarum (1753) [8] for botany, and the 10th edition of Systema Naturae (1758) [7] for zoology. Both publications include a standard naming system, a hierarchical classification system, taxonomic treatments reporting key characteristics and distribution, and citations of earlier publications. The value of drawings was not initially grasped and early works such as those of Linnaeus [7, 8] and the pioneer protistologists Otto Müller [19] lacked illustrations. The earliest illustrations recognized by zoological taxonomy appear in Aranei Svecici, a 1757 publication on the spiders of Sweden by C.A. Clerck [9]. Although actually published before the official start of zoological nomenclature, Aranei Svecici is explicitly recognized by the International Code of Zoological Nomenclature [20: Article 3]. Aranei Svecici contains illustrations of nearly 70 spider species. Nearly all of these feature a full body illustration (habitus) showing the dorsal view with legs symmetrically arranged. In a few cases, the male intromittent organ (the pedipalp) is illustrated. In many taxonomic groups including spiders, reproductive structures are rich in complex characters that show consistency within and differences between species [21, 22]. This makes reproductive structures valuable for recognizing and classifying many taxa, and they are frequently depicted in taxonomic treatments. Pedipalps are a pair of leg-like appendages that arise from the anterior part of the spider. As such, a pedipalp can be positioned and viewed in a limited number of cardinal orientations. When extended straight ahead and rotated in a transverse plane, four anatomically significant views are exposed in increments of 90°: dorsal, ventral, and two lateral views. In Aranei Svecici, the illustrations of the genitalia are less consistent than the habitus, but all have the pedipalp oriented along a cardinal anatomical axis. In the case of the green huntsman spider *Micrommata virescens*, both the habitus and male pedipalp are included (Fig. 2). The left pedipalp is illustrated from the left side (the retrolateral view; 180° from the prolateral view). A more recent guide to the spiders of Great Britain and Ireland [23] includes illustrations of both the habitus and pedipalp in the same orientation as Clerk first depicted in Aranei Svecici. Unsurprisingly, the more contemporary examples are more detailed and accurate, and the orientation of the images on a page may differ. Nonetheless, the selection of what to illustrate and how to orient it are unchanged despite the nearly 230 years separating these two publications. Comparative anatomy is a dominant organizing principle in taxonomic publications, regardless of the domain of life concerned. Figure 3 shows illustrations of Parnassia palustris flower anatomy from Linnaeus to the late 20th century. Key structural features are consistently visible across this time series. Similarly, a series of 18th and 19th century illustrations of the false chanterelle mushroom Hygrophoropsis aurantiaca show the same developmental stages and highlight the same anatomical details (Fig. 4).

**Fig. 2.**
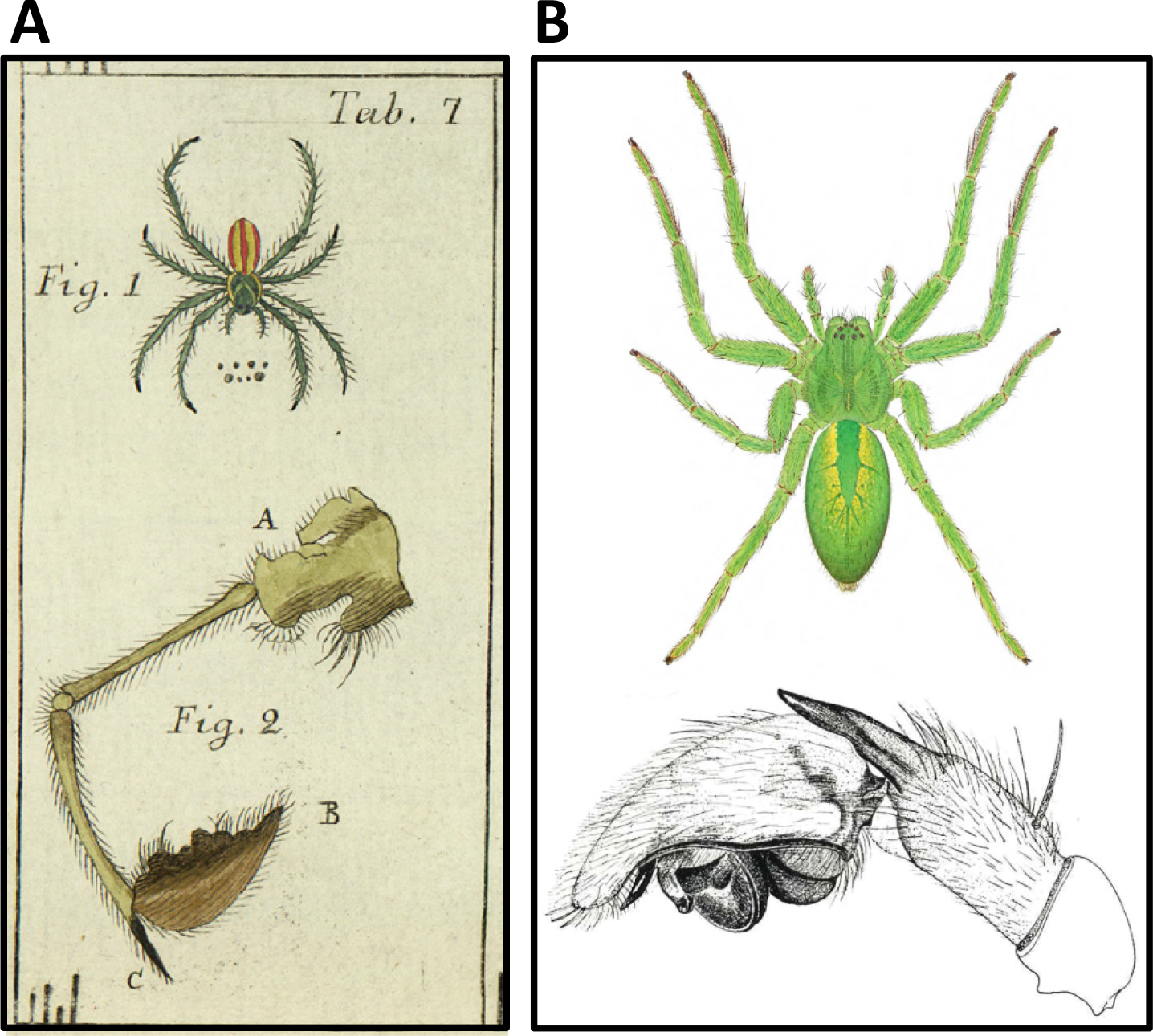
Time series of taxonomic illustrations depicting the spider *Micrommata virescens* (Arachnida: Araneae: Sparassidae) in standard views. (A) Illustrations from Clerck 1757 [9] (fig. 1, habitus, dorsal view; fig. 2, male pedipalp, retrolateral view). (B) Illustrations from Roberts 1985 [23] (top, habitus, dorsal view; bottom, male pedipalp, retrolateral view).

**Fig. 3.**
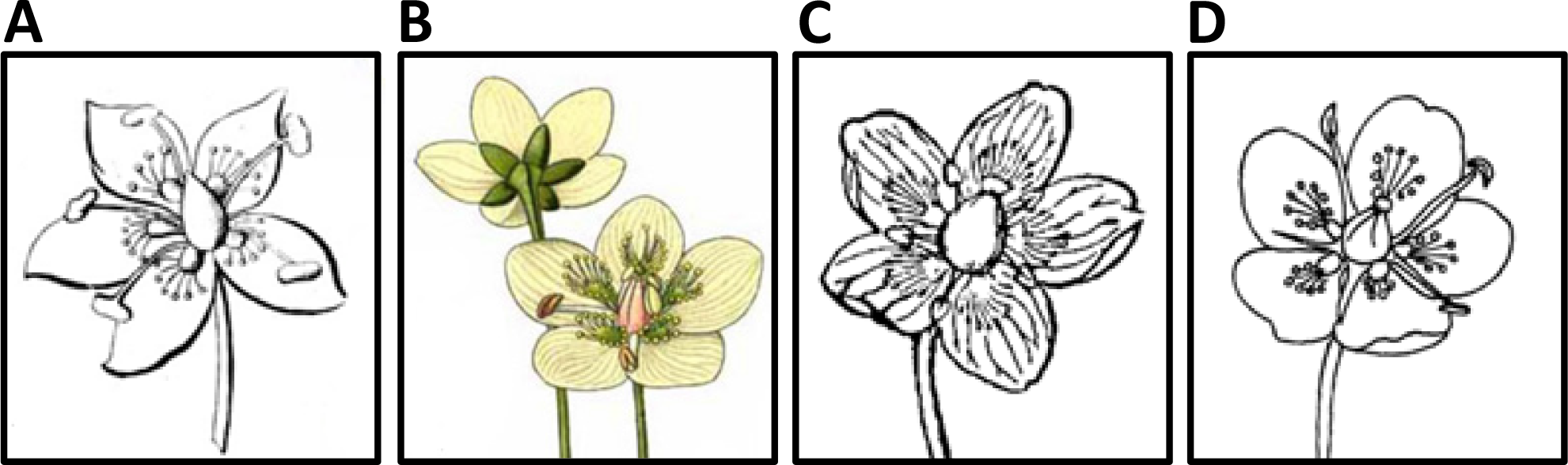
Taxonomic illustrations depicting the flower anatomy of the European marsh grass *Parnassia palustris* (Plantae: Angiosperms: Celastrales: Celastraceae). (A) From Linnaeus 1783 [24]; (B) From Masclef 1891 [25]; (C) From Britton and Brown 1913 [26]; (D) From Waterman 1978 [27].

**Fig. 4.**
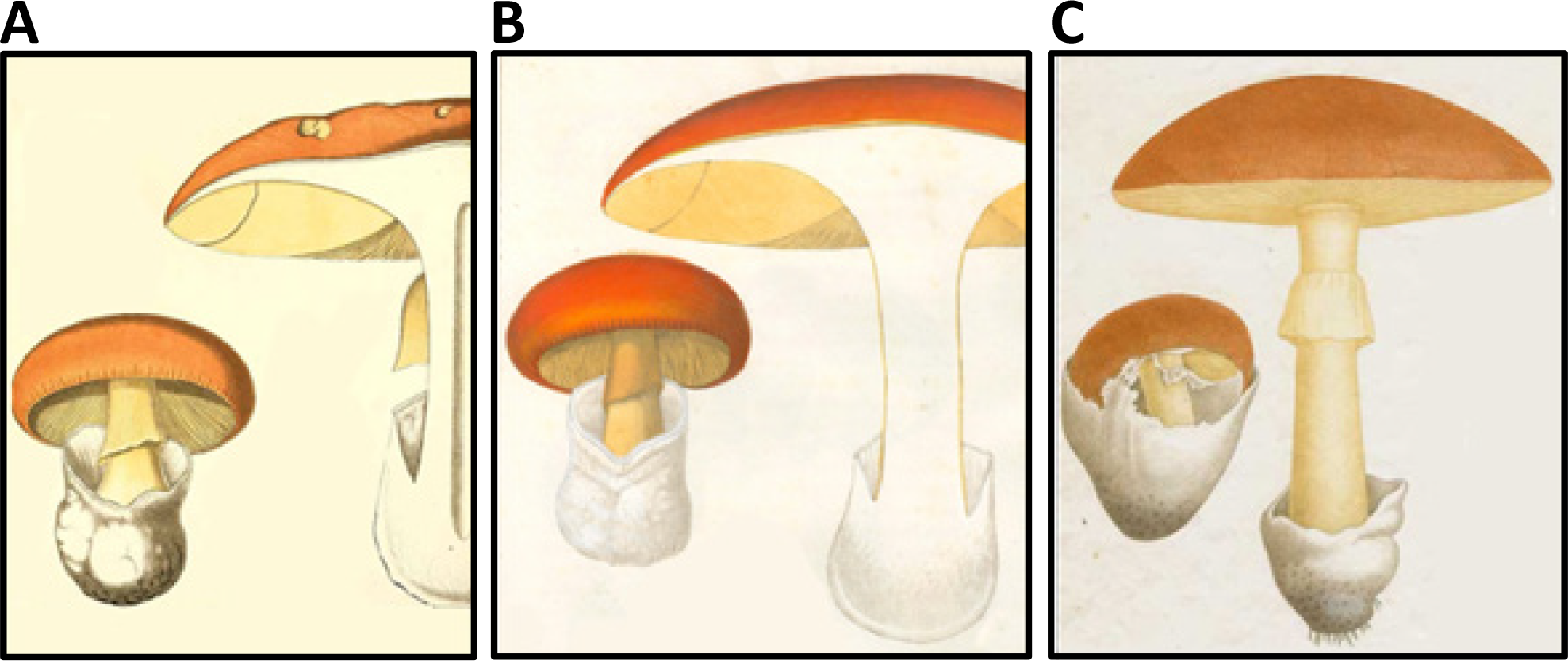
Taxonomic illustrations depicting the anatomy of the false chanterelle mushroom *Hygrophoropsis aurantiaca* (Fungi: Basidiomycota: Agaricomycetes: Boletales). (A) from Bulliard 1776 [28]; (B) from Bendiscioli 1827 [29]; (C) from Roques 1841 [30].

Foraminifera are single-celled amoeboid protists, mostly less than 1 mm in length, that typically construct a test (or shell). Although Foraminifera are relatively simple organisms, the cardinal orientations of their anatomy have long been recognized by taxonomists. Figure 5 includes excerpts from three taxonomic publications that deal with *Sigmolina sigmoidea.* It resembles a compressed sphere with a c-shaped pore at one end. A study from 1884 [31] and another from 1971 [32] depict this species with the same three standard views: a lateral view, a straight on view centered on the aperture, and an axial cross section. Another work from 1974 [33] depicts several *Sigmolina* species, but employs the same three standard views to depict and compare them. Despite the variety of forms in the axial cross section view of several species, the standard view makes them comparable.

**Fig. 5.**
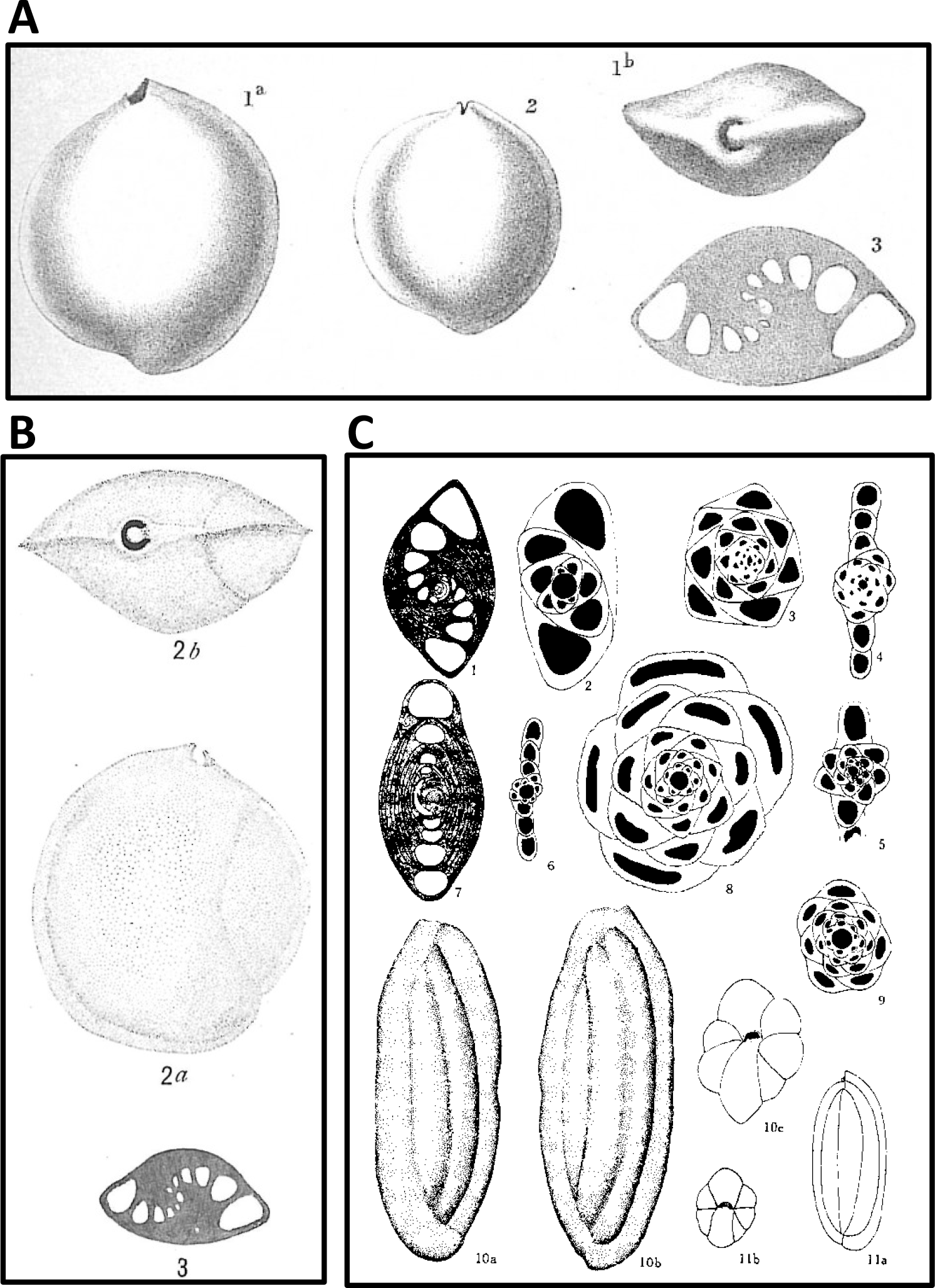
Time series of taxonomic illustrations depicting *Sigmoilina* (Chromista: Foraminifera: Miliolida: Hauerinidae) in standard views. (A) *Sigmoilina sigmoidea* from Brady 1884 [31] (1a, 2, lateral view; 1b, aperture view; 3 axial cross section). (B) *Sigmoilina sigmoidea* from Cushman 1971 [32] (2a, lateral view; 2b, aperture view; 3, axial cross section). (C) *Sigmoilina* species from Ponder 1974 [33] (1, *Sigmoilina sigmoidea*; 2-11, other *Sigmoilina* species; 1-9, axial cross section; 10a, 10b, 11a, lateral view; 10c, 11b, aperture view). A, B downloaded from World Register of Marine Species [34].

Scientific illustration can be expensive and time consuming to prepare, and costly to publish. This has historically placed limits on how thoroughly a treatment can be illustrated. For example, Biologia Centrali Americana (1879-1915) was a massive effort to document a regional fauna. It comprised 63 weighty volumes and included 1677 figure plates. But only 37% of the species treated were illustrated, and most of those species that were illustrated appeared in only one or two figures (Ramirez et al. 2007). Nevertheless, illustrations were generally limited to a few standard views. As in the previous examples, cardinal directions guide orientation. Figure 6 compares a plate from the first Biologia Centrali Americana volume on the insect order Orthoptera (grasshoppers, katydids, and their allies) [35] to Naskrecki’s more recent book on the Katydids of Costa Rica [36]. Both sources include a habitus in lateral view, habitus in dorsal view (which may be only partial), multiple views of the head region, and genitalia. Like most contemporary taxonomists, Naskrecki [36] depicts a core of standard views for all the taxa treated to facilitate comparison. The Orthoptera volumes of Biologia Centrali Americana depict many of the same standard views. But because many species are not illustrated for most standard views, there are gaps in knowledge that can make it difficult to apply Biologia Centrali Americana as a taxonomic guide.

**Fig. 6.**
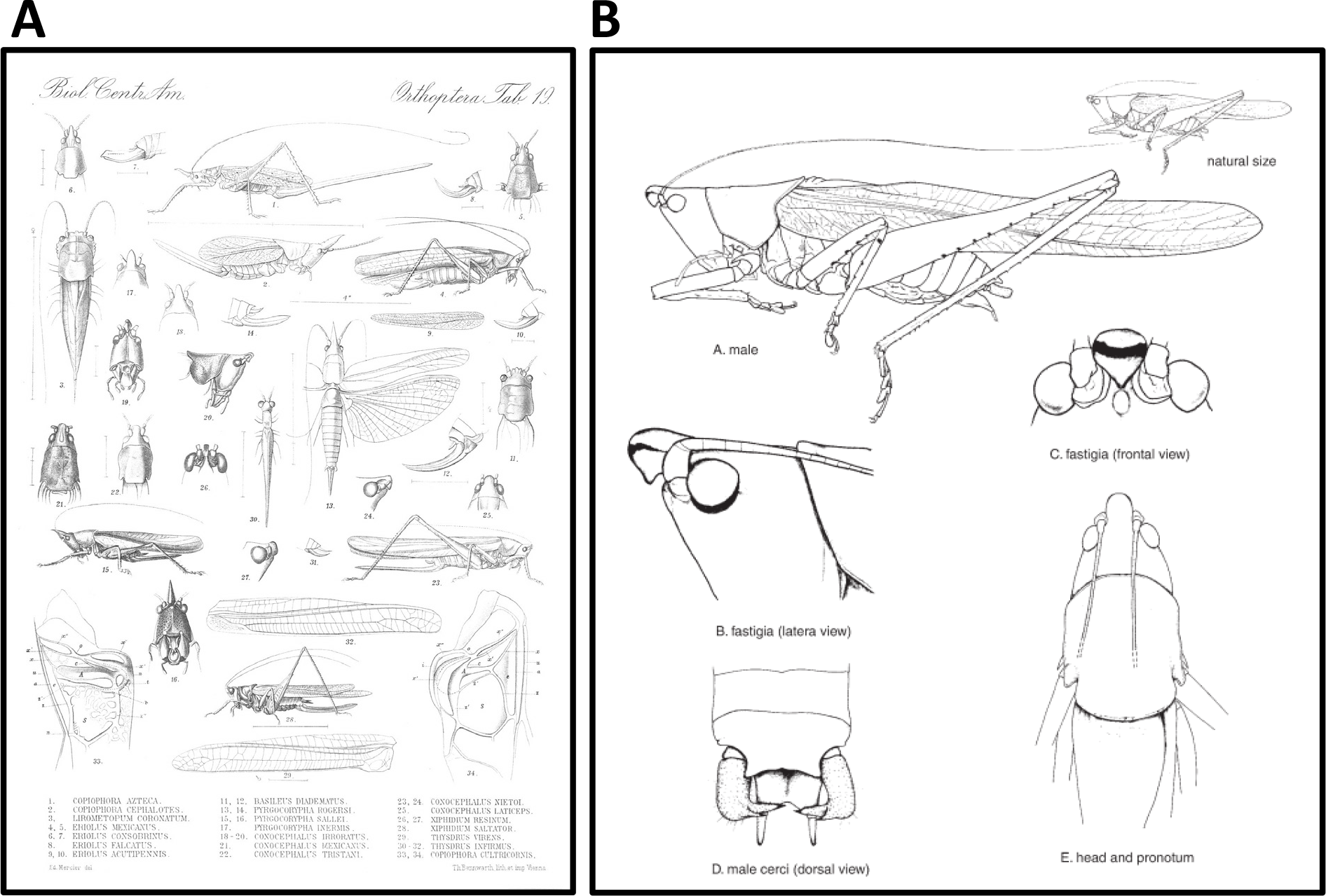
Time series of taxonomic illustrations depicting various katydid (bush crickets) species (Insecta: Orthoptera: Tettigoniidae) in standard views. (A) various conocephaline katydid species from Saussure 1898 [35], Plate 19 (1, 2, 4, 15, 23, 28, habitus of female, lateral view; 3, 13, habitus, dorsal view; 5, 6, 11, 17, 18, 21, 22, 25, 30, head region, dorsal view; 7, 8, 10, 12, 14, 31, female ovipositor, lateral view; 9, 29, 32, right forewing; 16, 19, 26, head region, frontal view; 20, 24, 27, head region, lateral view; 33, tambourine of left forewing, detail; 34, tambourine of right forewing, detail). (B) *Neoconocephalus affinis* from Naskrecki 2000 [36], fig. 12 (A, male habitus, lateral view; B, head region, lateral view; C, head region, frontal view; D, male cerci, dorsal view; E, head region, dorsal view. A accessed via Biodiversity Heritage Library (biodiversitylibrary.org/item/14636#page/484/mode/1up).

Interpretive difficulties arise when images of the same structure are not illustrated in the same way or from the same angle [37]. In an example from spider taxonomy, a 1942 publication by Chamberlin and Ivie [38] included treatments of nearly all known *Linyphantes* (Arachnida: Araneae: Linyphiidae) species, but did not include illustrations of the pedipalp in any commonly used orientation. The apical view is useful for distinguishing *Linyphantes* species from each other, but without also including widely used standard views, it is difficult to compare *Linyphantes* to other genera, such as *Bathyphantes*. In 1929, Petrunkevitch [39] published the only reference to include illustrations (albeit rudimentary) of both the retrolateral and apical views together (Fig. 7).

**Fig. 7.**
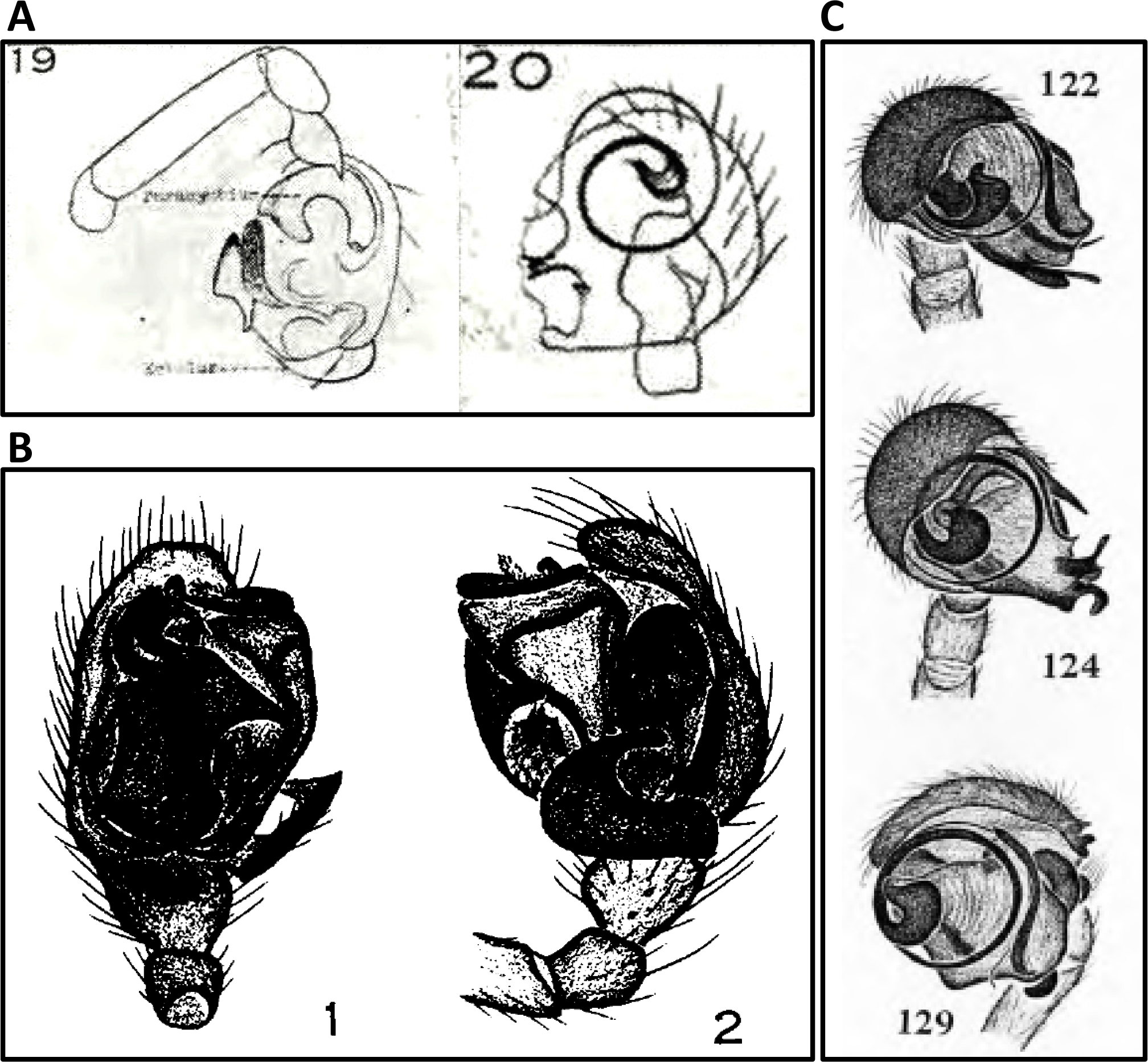
Taxonomic illustrations depicting illustrations of spider (Arachnida: Araneae: Linyphiidae) pedipalps from standard and non-standard views. (A) Illustrations of *Microneta aeronautica* (type species of genus *Linyphantes*, now called *Linyphantes aeronauticus*) from Petrunkevitch 1929 [39], Plate 1 (fig. 19, male pedipalp, standard retrolateral view; fig. 20, male pedipalp, rarely used apical view). (B) Illustrations of *Bathyphantes gracilis* from Ivie 1969 [40] (fig. 1, male pedipalp, standard ventral view; fig. 2, male pedipalp, standard retrolateral view); *Bathyphantes* may be a close relative of *Linyphantes*. (C) Illustrations of three *Linyphantes* species all from the rarely used apical view, from Chamberlin and Ivie 1942 [38].

As taxonomic knowledge within any particular group grows, community consensus about the relative value of various standard view images evolves. The importance of standard views to facilitate comparison has remained unchanged even as technologies and techniques have evolved, facilitating the inclusion of more numerous, high-quality images.

### 3.3 Forms of Images

Taxonomists and scientific illustrators use a variety of media to capture and convey the morphology and anatomy of organisms. Traditional techniques apply ink, graphite, paint, or other such substances alone or in combination to paper, board, or other such surfaces [15]. For most of the history of taxonomy, line drawings (black ink on paper) have been the most widely used medium for depicting anatomy, complemented by colored plates and, in the early 20th century, photography.

Starting around the mid-1980s, new technologies introduced alternative mechanisms for capturing and rendering information about morphology. Computer-aided illustration techniques were developed. Mixed media approaches made it possible to combine multiple techniques into single composite images, such as a body rendered by hand in pencil combined with photographs of wings (Fig. 8A). The increasing availability of scanning electron microscopy (Fig. 8B) opened new frontiers of discovery [41-43]. Advances in digital cameras mounted on microscopes, and the advent of extended focus composite imaging [44, 45] reduced the time and expense of graphically representing morphology (compared to illustration in particular), and photographs began to eclipse illustrations as the primary means of documenting morphological structures (Fig. 9; e.g., Riedel et al. 2013 [46]).

**Fig. 8.**
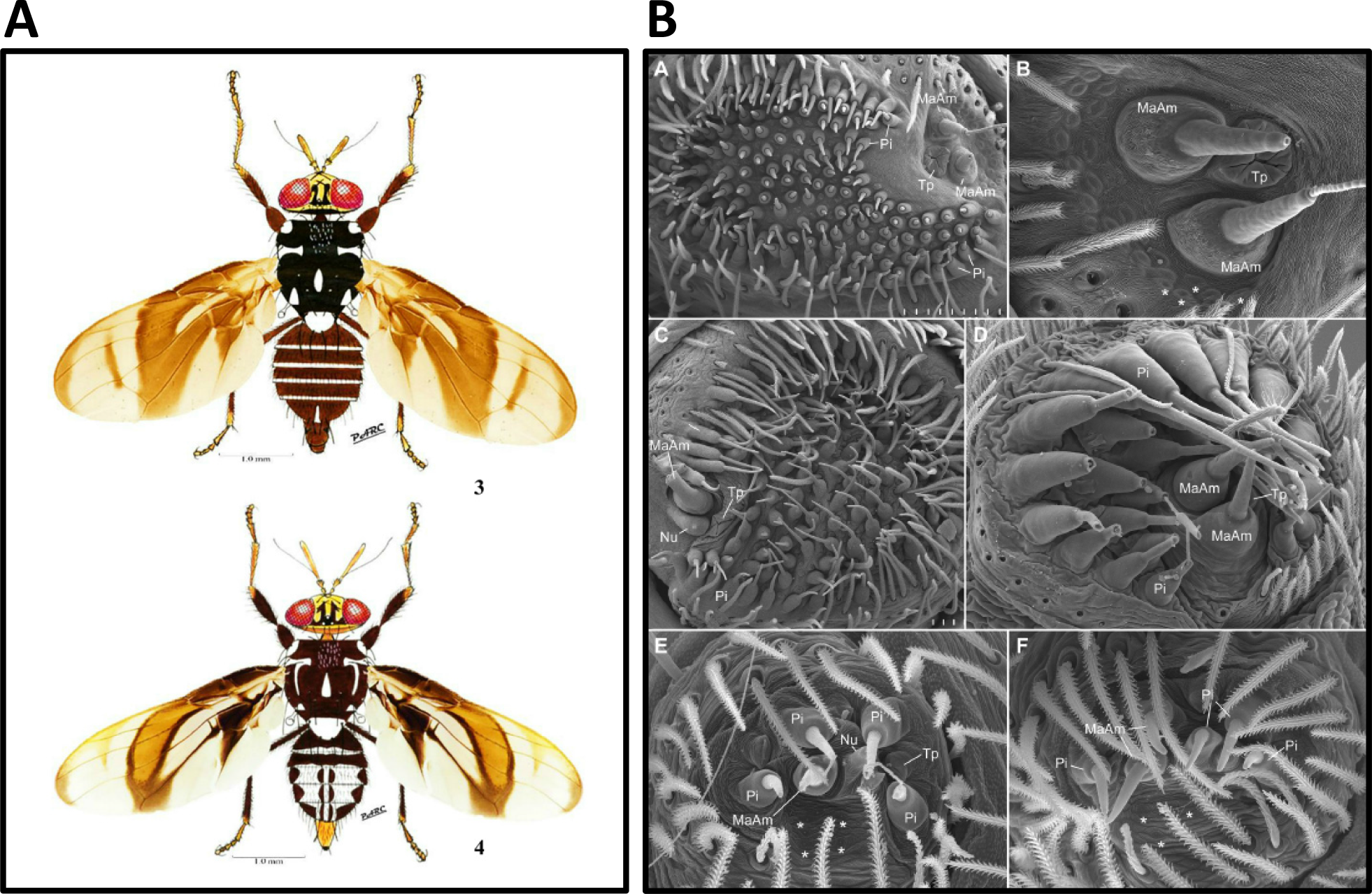
Use of alternative media to depict and compare anatomy. (A) Mixed media representation of two fly species. Wings are photographs while other parts were illustrated with color pencils. from Rodriguez et al. 2016 [47] (fig. 3, *Cryptodacus ornatus*; fig. 4, *Cryptodacus trinotatus*). (B) Scanning electron microscope images comparing the spinnerets of various spider species, from Ramírez et al. 2014 [48] (anterior lateral spinnerets, E, C, male, others female; A, B, Austrochilidae: *Thaida pecularis*; C, Tengellidae: *Tengella radiata*; D, Homalonychidae: *Homalonychus theologius*; E, F, Penestomidae: *Penestomus egazini*).

**Fig. 9.**
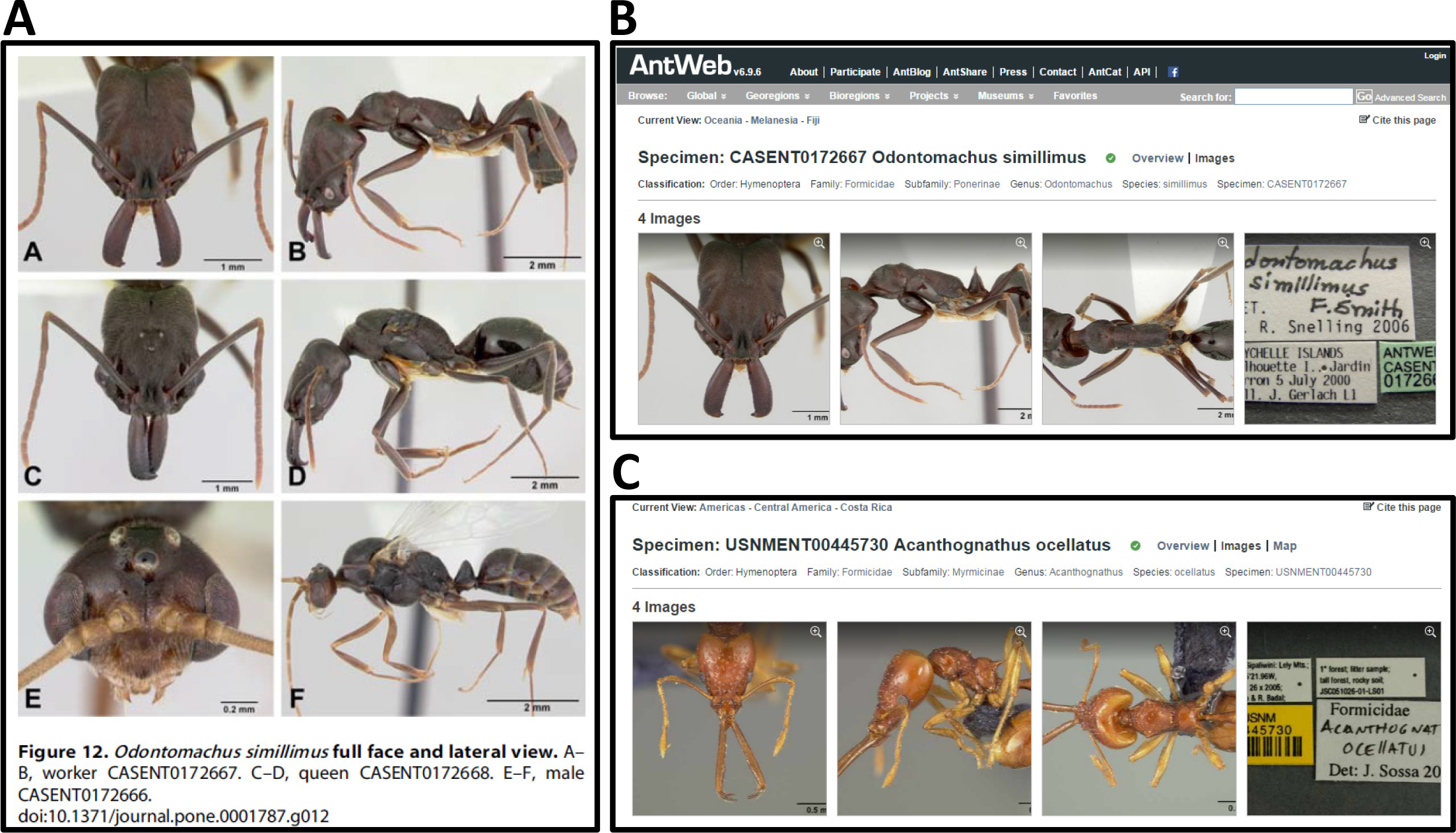
Extended focus composite photographs of ants in a taxonomic publication and the AntWeb online database. (A) Head and profile views of three specimens of the ant *Odotomachus simillimus*, from Fisher and Smith 2008 [49]. (B) the ant *Odontomachus simillimus* on AntWeb, same specimen as top row in A. (C) the ant *Acanthognathus ocellatus*. B and C were contributed by different research labs both following AntWeb’s imaging protocol to facilitate comparison.

Other radiation imaging techniques, such as X-rays, are used to detail skeletal elements in animals, and tomography (micro-CT, synchrotron) is increasingly used to compare detailed anatomy with the aid of three dimensional computer models (Fig. 10; [50]). With three dimensional interactivity, structures can be compared from any angle. Sonograms are used to represent and compare the sounds made by organisms such as birds, crickets, bats, and whales (Fig. 11).

**Fig. 10.**
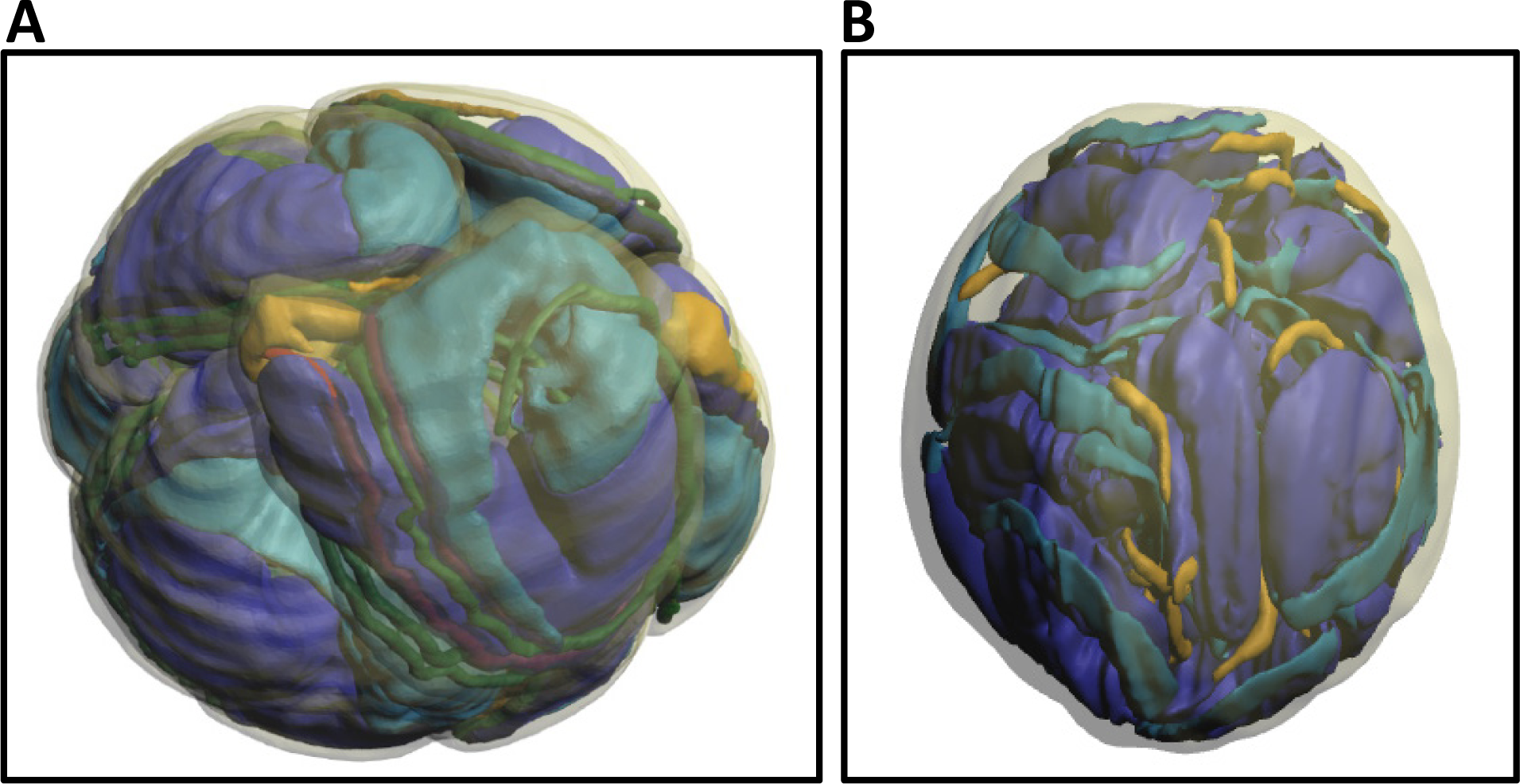
Surface renderings of spider sperm reconstructed based on digital tomography. (A) *Kukulcania hibernalis* (Filistatidae), from Michalik and Ramírez 2014 [51], with credit to E. Lipke. (B) *Orsolobus pucara* (Orsolobidae), from Lipke et al. 2014 [52]. The PDF file of this article contains interactive 3D content. Click on the image to activate content and use the mouse to rotate objects. Additional functions are available through the menu in the activated figure.

**Fig. 11.**
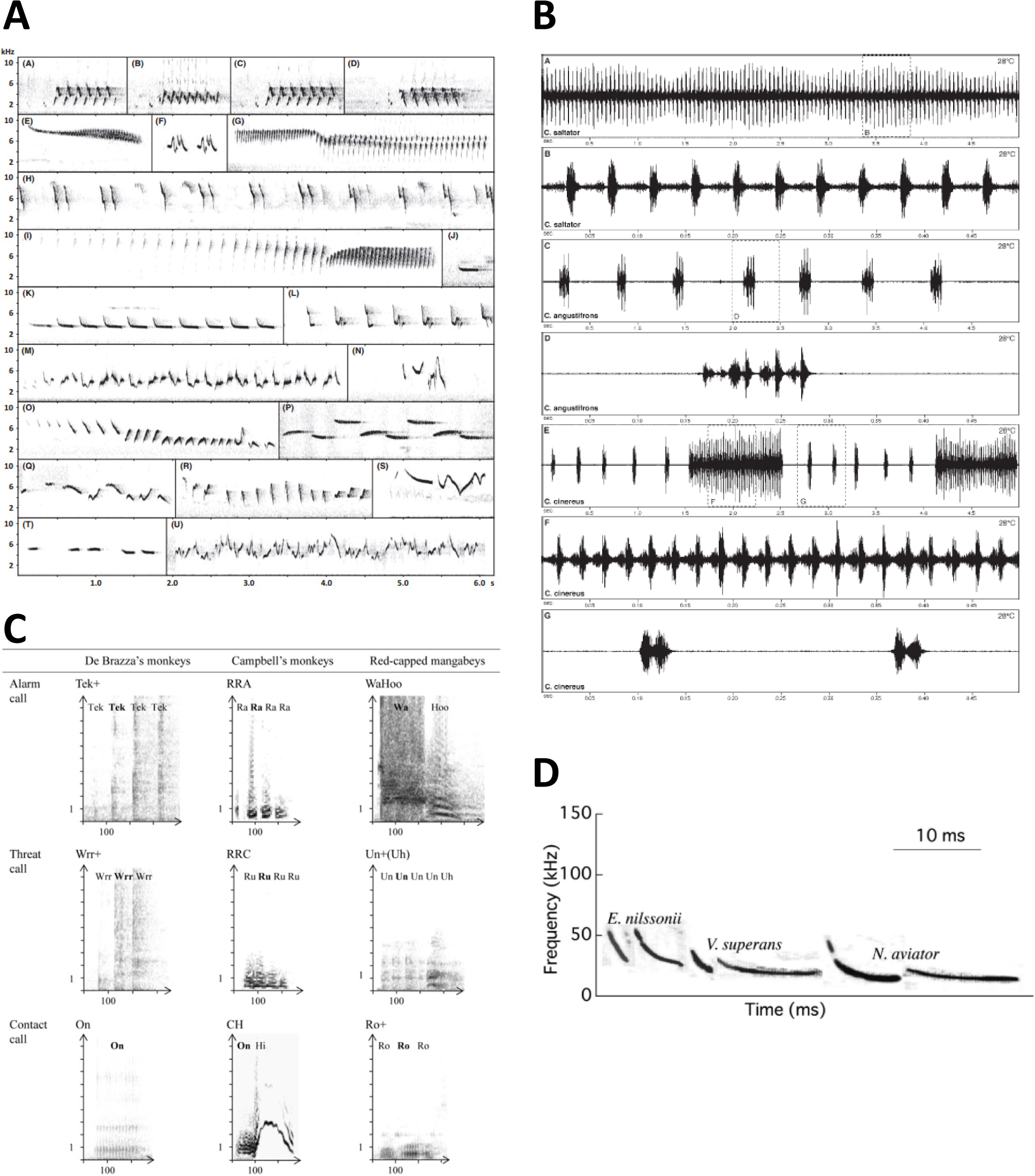
Comparative sonograms visualizing sounds made by a selection of animal groups. (A) Songs of assorted leaf warbler species (Aves: Passeriformes: Phyllosopidae: *Phylloscopus*), from Tietze et al. 2015 [53]. (B) Oscillograms showing two types of male airborne calls from three species of katydid (Insecta: Orthoptera: Tettigoniidae: *Conocephalus*), from Naskrecki 2000 [36]. (C) Three different call types (alarm, threat, and contact) across three monkey species (Mammalia: Primates: Cercopithecidae), from Bouchet et al. 2013 [54]. (D) Echolocation calls of three bat species, two of each included to show some intraspecific variation (Mammalia: Chiroptera: Vespertilionidae), from Fukui et al. 2004 [55].

Taxonomic publications often feature photographs from the field, typically depicting living organisms and their habitats (Fig. 12). Such photographs may not be structured and standardized with the precision of a standard view anatomical illustration, but the purpose of such photographs is to document facts, such as color and posture of the organism in life, habitats where it has been found, behavior and interactions with others, and more. Aesthetic and artistic considerations are secondary.

**Fig. 12.**
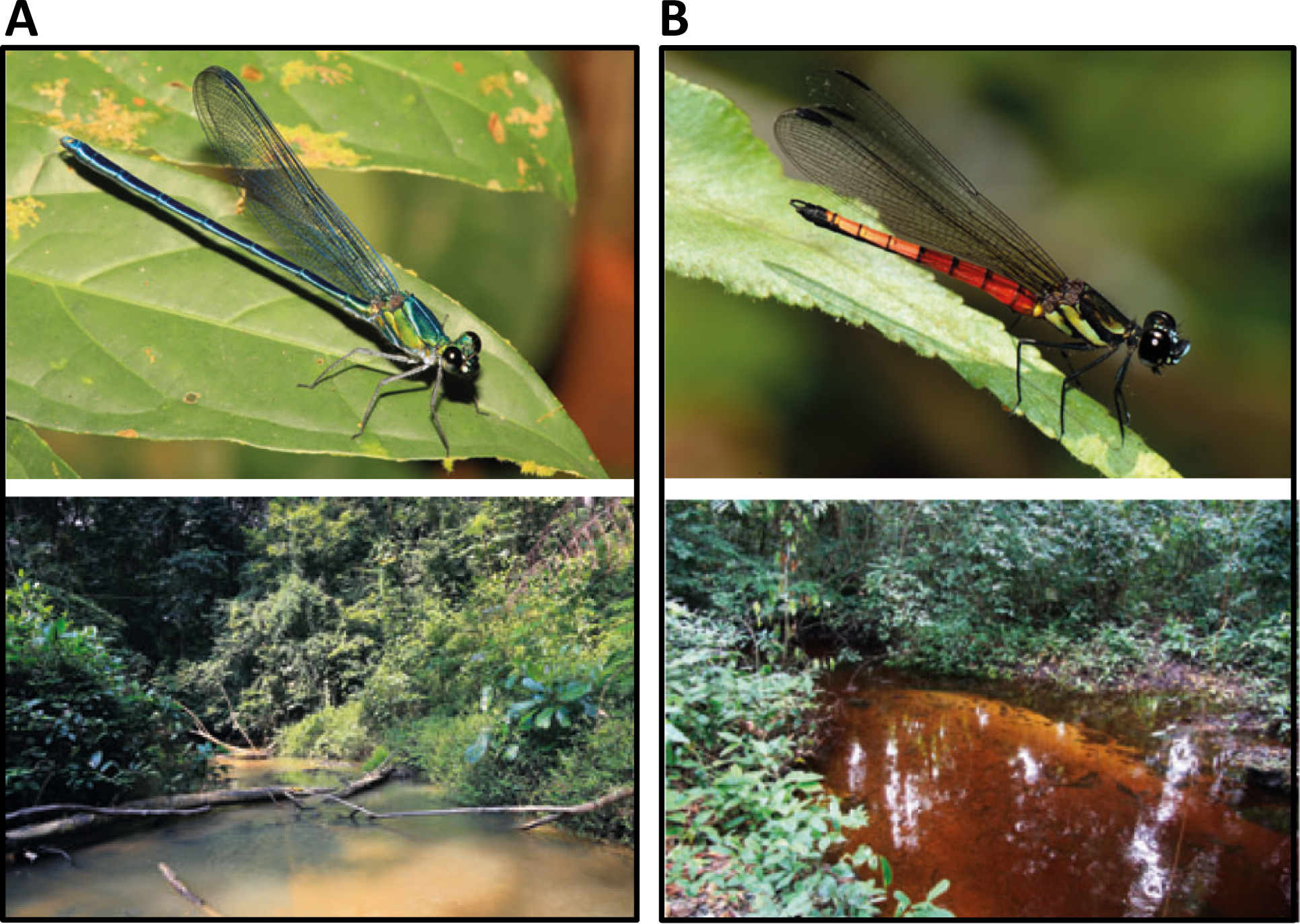
Semi-standardized photographs depicting live animals in the field and associated habitats. A, the damselfly *Umma gumma* (Insecta: Odonata: Calopterygidae), male specimen and habitat. B, the damselfly *Africocypha varicolor* (Chlorocyphidae), male specimen and type locality. From Dijkstra et al. 2015 [56].

Various automated methods are increasing used to capture information. Camera traps are automated image capture systems left in the field for an extended time to document wildlife activity in a particular location [57] (Fig. 13). Camera trap images rarely feature in true taxonomic publications, but contribute knowledge about where a particular species occurs, and thus provide observation data for scientific publications and conservation management [58-60]. Other automated techniques included mass-digitization of museum and herbarium specimens [16, 17, 61, 62] (Fig. 14), robotic imaging of the sea floor or other inaccessible habitats [60, 63], and flow cytometers with automatic image capture to take pictures of phytoplankton [64, 65].

**Fig. 13.**
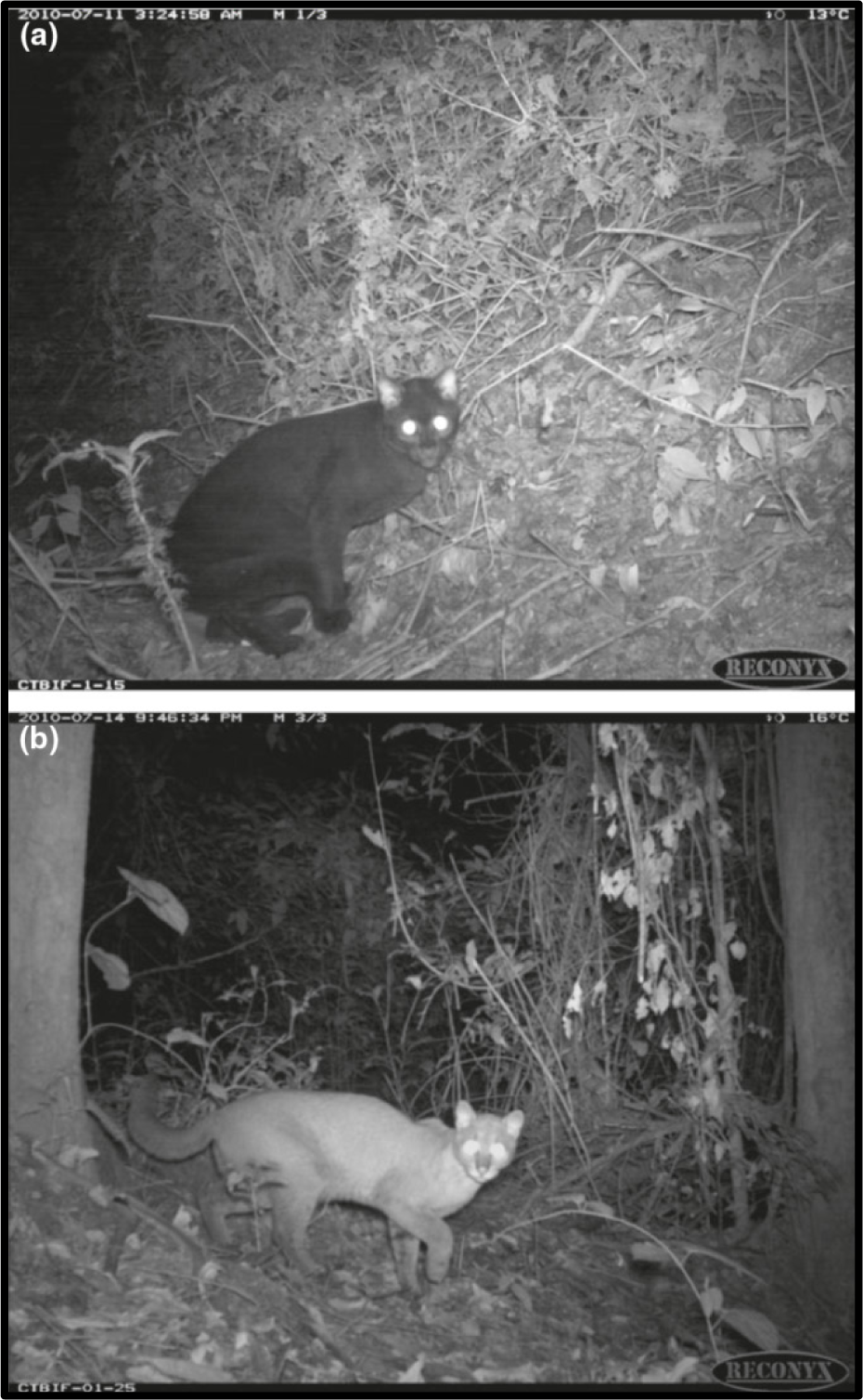
Camera traps document species occurrence. African Golden Cat (Mammalia: Carnivora: Felidae: *Caracal aurata*, formerly called *Profelis aurata*) in Bwindi Impenetrable National Park, Uganda (A, dark color morph; B, light color morph). From Mugerwa et al. 2012 [66].

**Fig. 14.**
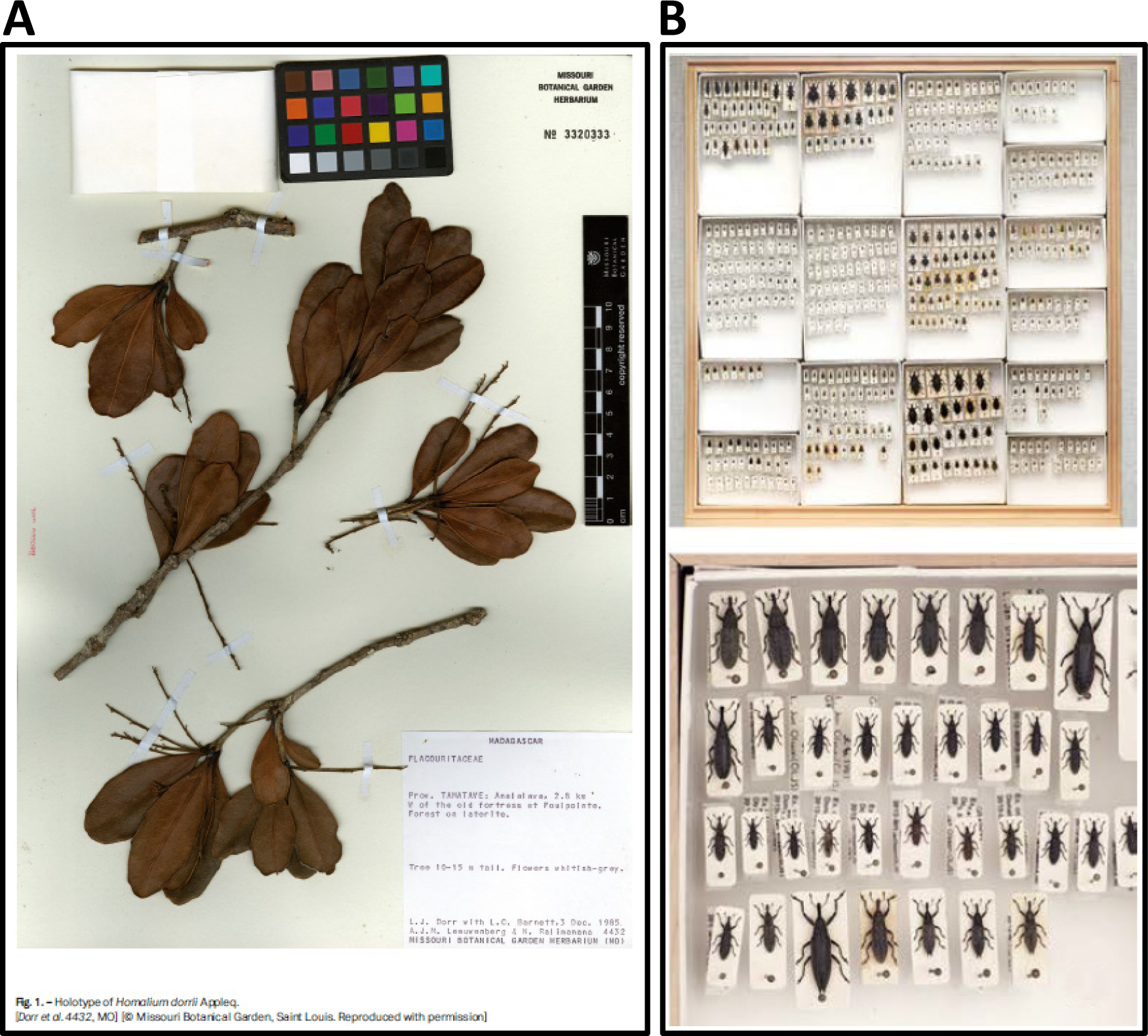
Images of specimens from museum collections. (A) Herbarium sheet of a holotype specimen (Angiosperms: Malphighiales: Salicaceae: *Homalium dorrii* Appleq.), specimen 3320333 of the Missouri Botanical Garden, from Applequist 2015 [67]. This is one of many thousands of herbarium sheets digitized by a semi-automated process at herbaria worldwide. Note the copyright declaration on the scale and in the original figure caption. (B) Entire entomological collection drawer imaged using high resolution semi-automated method. Lower image is detail from upper left corner of drawer, from Holovachov et al. 2014 [61].

Of special significance to taxonomists are images of specimens that are held in institutions such as museums and herbaria. These are estimated to be over 3 billion specimens in about 55,000 museums and 3,500 herbaria around the world. Many are of individual organisms that are significant to the taxonomic or nomenclatural history of the taxon. Of these, the most important are the organisms that are type material as they help establish the identity of taxa. All other specimens help to clarify variation within and among species. As taxonomists need to inspect the materials on which nomenclatural and taxonomic decisions are made, they require access to the preserved material. Historically, taxonomists had to visit repositories or have materials shipped to them. This was costly and specimens were at high risk of being damaged if shipped around the world. Now the use of high resolution images inclusive of 3D images is effective for most purposes, cheap and with low risks of damage. This has led to the investment in specimen digitization programs, such as iDigBio is the US [16]. Taxonomic materials are presented using standard techniques, such as pinned insects or herbarium sheets for plants [68-72].

Taxonomic publications often include maps, typically to show the geographic distribution of occurrence records. Base maps may be static, from a printed or graphical source, or rendered as a layer in a GIS (Geographic Information System) environment. Google allows annotation and non-commercial distribution of its maps including their use in journal articles when proper attribution is provided (Fig. 15; google.com/permissions/geoguidelines.html).

**Fig. 15.**
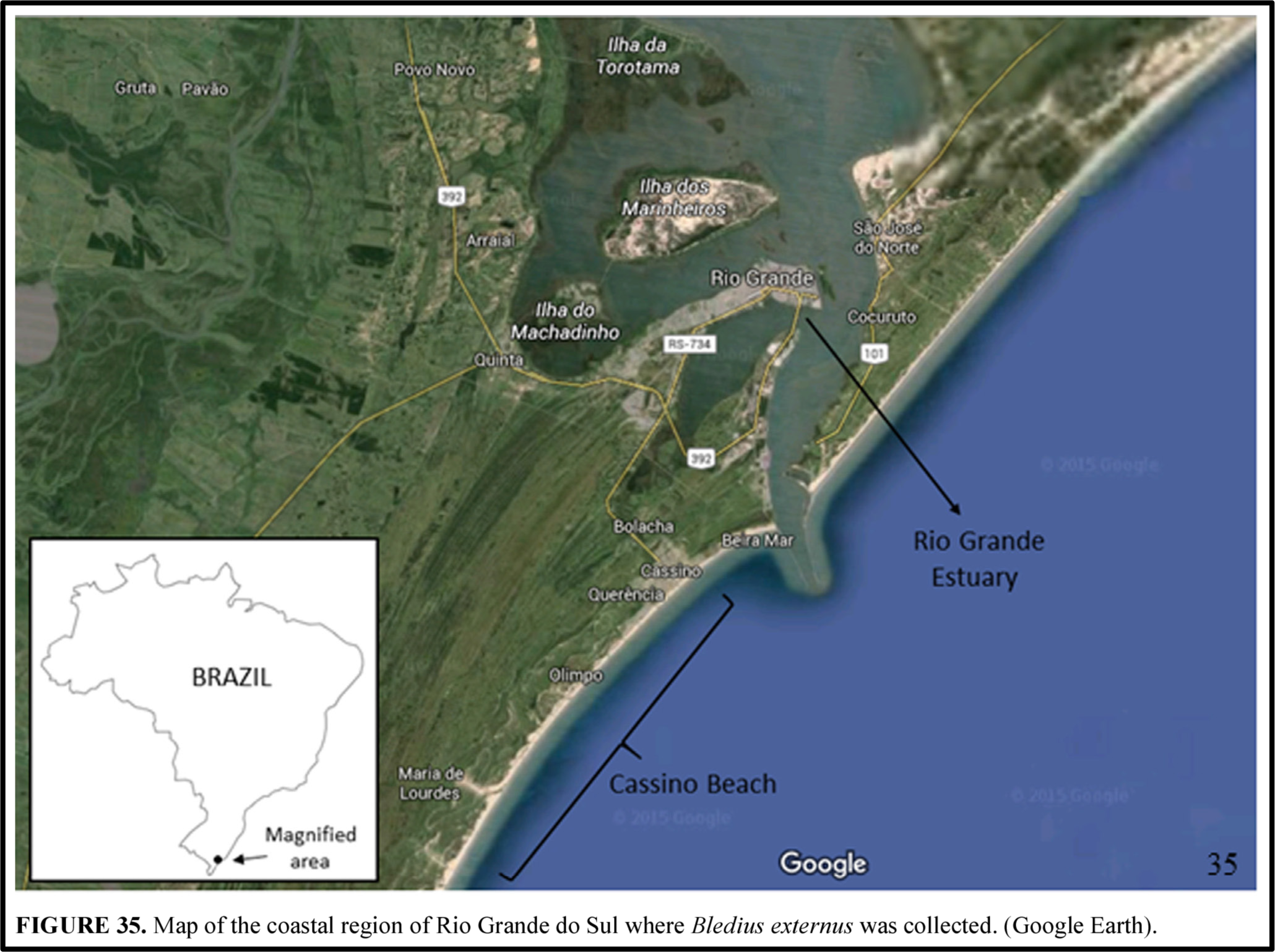
Composite map showing region where the beetle *Bledius externus* (Insecta: Coleoptera: Staphylinidae: Oxytelinae) was collected. This map incorporates elements obtained from Google Earth attributed to their source. From Castro et al. 2016 [73].

Individual figures are often combined to form plates composed of several species to facilitate comparison (Fig. 16). The images may be arranged to represent relative size (with larger and smaller subjects shown at a common scale), or at different scales with a scale bar included to insure that actual size can be inferred. Such plates are especially common in field guides, where the primary purpose is efficient taxonomic determination.

**Fig. 16.**
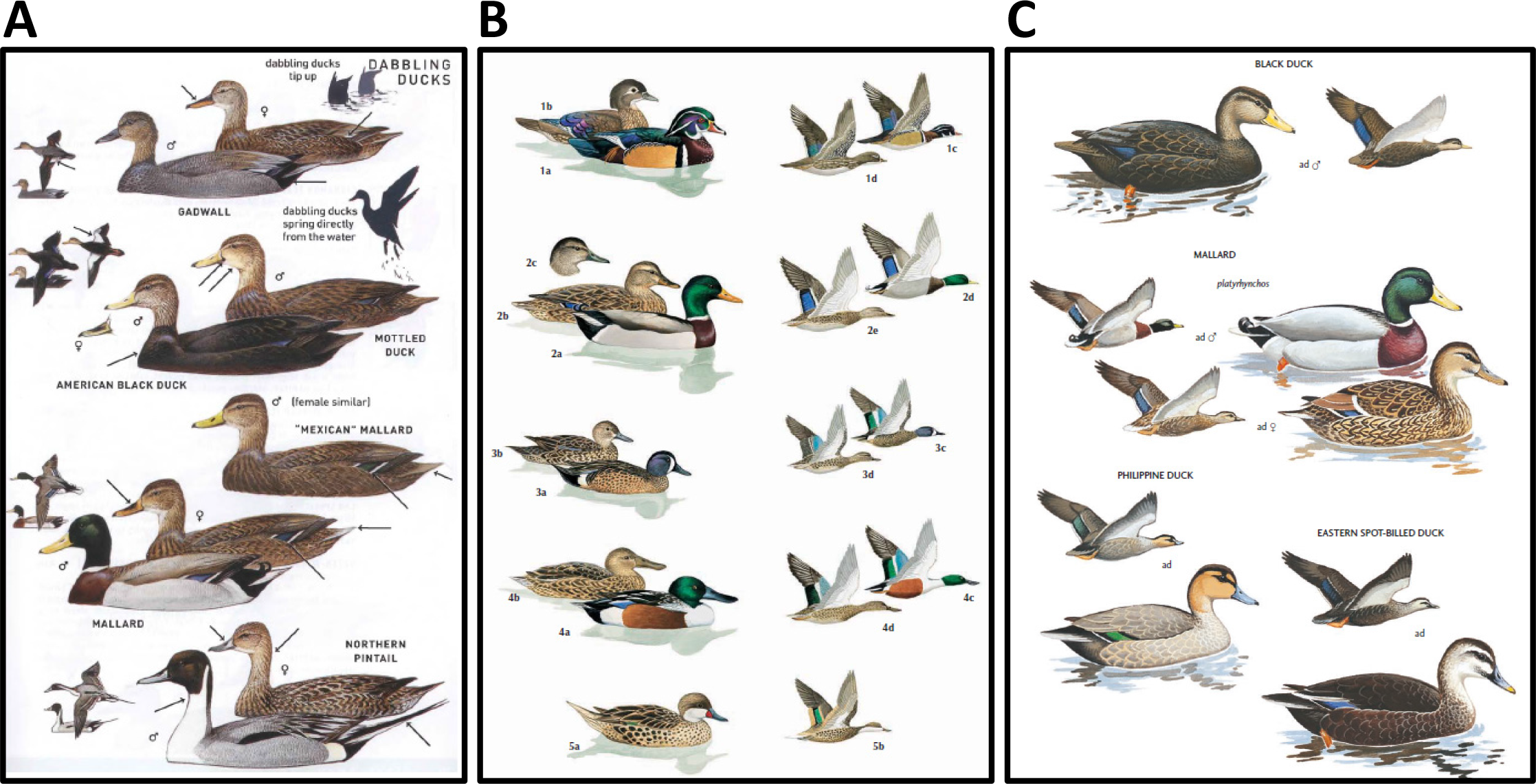
Color plates from field guides to birds (Aves). Note repeated depictions of different sexes and behaviors. (A) from Peterson 2008 [74]. (B) from Latta et al. 2006 [75]. (C) from Brazil 2009 [76].

Quantitative data may be represented as scatter plots (with or without trend-lines), graphs, histograms, pie charts, and other such devices. Charts provided for the purpose of establishing or comparing the distinguishing characteristics of a species lack the requisite creative element that makes copyright applicable (Fig. 17).

**Fig. 17.**
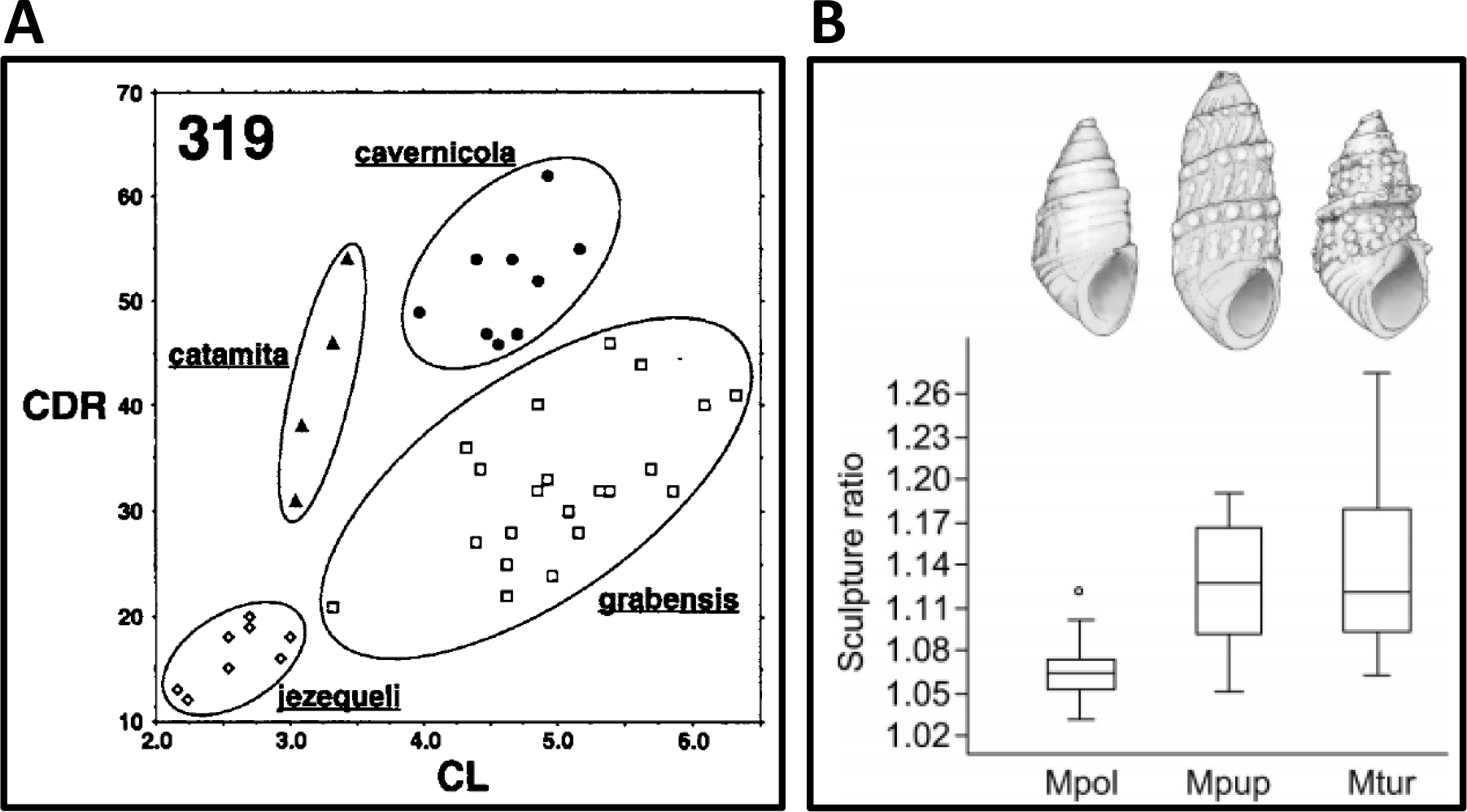
Visualizations of diagnostic morphometric characters. Quantitative characters, alone or in combination, can contribute to taxonomic identification. Values from an unknown specimen can be compared to those presented in charts such as these. (A) Scatter plot of two morphometric values for four spider species (Araneae: Dipluridae: *Lathrothele*), each with a distinct domain, from Coyle 1995 [77]. (B) Sculpture ratio, a quantification of shell texture based on a ratio of two measurements, for three Holocene snail species (Mollusca: Gastropoda: Thiaridae: *Melanoides*), from Bocxlaer and Schultheiß 2010 [78].

The criteria for determining whether copyright applies to a class of images are the same regardless of subject matter. We emphasize here those classes of images most applicable to taxonomy, but the principle applies to other domains of science. That is, if the image adopts conventions intended to facilitate comparison with other works, then the image is unlikely to be a creative work in the sense of copyright. This does not mean that images in taxonomy are less important than those from creative fields, only that copyright protection is neither applicable legally nor desirable in the context of comparative science.

## 4. Rights and scientific images

It is a widespread belief among biologists that scientific images are “owned” by somebody, such as the author, photographer, the institution that employs the creator, or the publishing house responsible for publishing the images [12]. The notion of “ownership” carries with it a sense of ownership akin to that applied to tangible goods. This may lead to the presumption that property rights apply. Such rights may be used to assess a monetary value, limit access, and prescribe how goods may or may not be reused by others. Every physical thing can by default be an object of property. But property rights apply only to tangible goods. There are exceptions or limitations to property rights. These exceptions and limitations are defined by national laws and can vary slightly from country to country. Such exceptions may refer to out-of-commerce goods as air, water, mountains with the exceptions dating back to Roman Law where they qualified as *“res communes omnium”.* Another suite of exceptions are based on ethical reasons and include, as an example, the physical integrity of individuals.

Imbuing images with a sense of ownership as if they were tangible goods is misleading, because images are not tangible goods [79]. Taxonomists do not perceive the value of a biological illustration as arising from the original physical ink on paper (or other media), nor in terms of its artistic appeal and distinctiveness, but rather from the concept or insight that is depicted. A concept or insight is ‘intellectual property’ and is not a tangible good. That is to say, only rights relating to non-tangible goods are relevant here. So for legal issues related to images, we must look among the rules applicable to rights in non-tangible goods - that is, to intellectual property rights. These are based on different principles than those for tangible property rights. In the following sections we discuss the bases of intellectual property rights in creative works and how they differ from ownership rights.

### 4.1 Numerus clausus of Intellectual Property Rights

National laws specify which non-tangible goods may be regarded as intellectual property. As is the case with property rights with respect to tangible goods, each country may have different rules for intellectual property rights. International instruments such as treaties and conventions aim to harmonize national rules and reduce discrepancies by fixing minimum standards and recommending rules for the application of rights. Various international instruments address specific branches of intellectual property. Well known examples include the Berne Convention for the Protection of Literary and Artistic Works (http://www.wipo.int/wipolex/en/details.jsp?id=12214), the Rome Convention for the Protection of Performers, Producers of Phonograms and Broadcasting Organizations (http://www.wipo.int/wipolex/en/details.jsp?id=12656), and the Paris Convention for the Protection of Industrial Property (http://www.wipo.int/wipolex/en/details.jsp?id=12633). These international instruments apply only to the extent that they are represented in national laws.

The protection of non-tangible goods is always limited to specific areas and specific objects; there is no “general protection”. If national laws do not specify that particular non-tangible goods are objects of intellectual property rights, then no rights apply. Individuals cannot claim intellectual property rights over items that are not covered by the relevant national laws.

Intellectual property rights with respect to non-tangible goods is always limited to a restricted number (the so-called “numerus clausus”) of specifically attributed rights [80]. With regard to scientific images, there are only four relevant areas of intellectual property rights: copyright (or “authors’ rights”, as it is referred to in international conventions and in most European countries), the EU-specific database protection, protection against unfair competition, and, in a few European countries, special protection for photographs. We discuss these areas below.

### 4.2 Copyright

Copyright protects certain human creations in art and literature. The minimum standard of this protection, applicable within the 172 countries that have introduced this form of intellectual property rights into their national laws, is defined in the Berne Convention for the Protection of Literary and Artistic works. The Convention was first established in 1886 and has been amended several times (http://www.wipo.int/wipolex/en/details.jsp?id=12214).

Article 2 paragraph 1 of this Convention declares that “the expression ‘literary and artistic works’ shall include every production in the literary, scientific and artistic domain, whatever may be the mode or form of its expression, such as books, pamphlets and other writings; (…) works of drawing, painting, architecture, sculpture, engraving and lithography; photographic works to which are assimilated works expressed by a process analogous to photography (…).” Member countries of the Berne Convention are therefore obliged to protect illustrations such as “drawings” and other artistic works or “photographic works” by their national copyright law [81].

Copyright confers to the author a set of privileges which result in far-reaching control over access to the work and over most forms of re-utilization. These rights are limited in time (normally to 50 or 70 years after the author’s death) and may be restricted by numerous legally defined exceptions and limitations. Again, there are important differences from country to country with respect to the content and the extension of rights conferred to authors as well as to the definitions of exceptions and limitations. In the United States of America, the “fair-use-principle” (see below 5.3) substitutes for the concept of exceptions and limitations.

When a concept or intent is captured in the form of a photograph, graph, drawing or illustration, it is said to be ‘fixed’ or ‘expressed’. The Berne Convention (Art. 2 par. 2) allows the member countries “to prescribe that works in general or in any specified categories of works shall not be protected unless they have been fixed in some material form”. This restriction exists in the United States of America and in many other jurisdictions.

The Berne Convention does not define the notion of “work”, but leaves that definition to national legislators. As a consequence, the notion of “artistic work” or “photographic work” can vary from country to country. However, there are aspects of this term that are identical in all copyright systems. One of them is that “works” must be man-made. Objects produced by nature or by organisms never qualify as copyrightable. Another important criterion is that intellectual productions qualify as works only if they are somehow original. This criterion does not refer to the content of the work, but to the form of presentation [81, section 2.8]. Copyright applies to a “work” only if it is expressed in an original (individual, new, creative) way [82]. In the case of a photographic picture, it can only be considered as a copyrightable photographic work if it somehow differs in style compared to other photographs taken of the same object or other similar photographs with which it may be compared.

The same applies to drawings and other forms of scientific illustrations: they are works in the sense of copyright if they adopt an original form of expression. Illustrations that follow predefined rules or conventions do not qualify as copyrightable works. Illustrations of biological information, especially in taxonomy, usually follow conventions that facilitate comparisons with similar illustrations. When this is the case, the images do not qualify as copyrightable works.

According to U.S. Copyright Law, a work may not qualify for copyright protection if it is about an “idea, procedure, process, system, method of operation, concept, principle, or discovery, regardless of the form in which it is described, explained, illustrated, or embodied in such work.” 17 U.S. Code § 102 (b) https://www.law.cornell.edu/uscode/text/17/102. The paragraph describes a concept that is basic for all copyright laws in the world [81, section 2.8].

The mechanical reproduction of an already existing photograph, drawing, painting or other forms of two-dimensional presentation (such as herbarium sheets) cannot qualify as photographic works for copyright purposes [83]. Objects of photographic works must be three-dimensional. If the two-dimensional object of the photograph is a copyrightable work, the photographs qualify as reproductions of the copyrighted work, but are not photographic works in themselves.

Copyright protection, does not only refer to single works, but also to collections of objects (Art. 2 par. 5, Berne Convention). Again, the qualification as copyrightable work requires that there is originality and individuality in the selection or in the arrangement of the objects. As established by the U.S. Supreme Court decision in Feist Publications, Inc. v. Rural Telephone Service Co. (499 U. S. 340, 1991), collections of objects that are presented in a standardized form, for example in alphabetical order, or in the case of Biology following a predefined systematic order, will not qualify as copyrightable works. The inclusion of single drawings within a plate to combine, summarize or compare attributes of organisms also follows established conventions and such plates are therefore ineligible as copyrightable works.

### 4.3 Database protection

In most countries, databases are protected by intellectual property rights to the extent that they qualify as works in the sense of copyright. This is the case where there is individuality in the selection of data or in the form of presentation of these data. Databases that do not meet these requirements are not subject to specific protection rules.

An important exception to this rule exists in the E.U. The E.U. introduced, with directive 96/9/EC (http://eur-lex.europa.eu/legal-content/EN/TXT/PDF/?uri=CELEX:31996L0009&rid=1), a special protection for databases that is independent of, and in certain cases complementary to, copyright protection. This so-called *sui generis* protection applies to databases which show “that there has been quantitatively and/or qualitatively a substantial investment in either the obtaining, verification or presentation of the contents” (art. 7, Directive 96/9/EC). This allows the person who invested in the creation of the database to prevent the extraction or re-utilisation of the whole or a substantial part of the contents of that database.

The term “database” is defined in Directive 96/9/EC art. 1 no. 2 of the directive: “For the purposes of this Directive, ‘database` shall mean a collection of independent works, data or other materials arranged in a systematic or methodical way and individually accessible by electronic or other means.” This is consistent with a data environment being structured into one or more tables, of tables having one or more fields, and fields holding data. The fields are defined by metadata. The content of such databases is made visible or can be copied using web-services or web interfaces.

Database protection does not deal with individual data elements. The intellectual property right refers to the database as a whole, not to an individual datum. Database protection may therefore apply to a collection of scientific images, but not to an individual image. The protection is very specific to prevent the extraction and reutilization of the database as a whole or of substantial parts of it. It does not serve to prevent the extraction and re-utilization of individual data or of groups of datasets that do not constitute a substantial part of a database.

As the European Court of Justice has pointed out in several judgments, European database protection concerns the creation of databases out of material that already exists, but does not deal with the creation of data as such. “Investment in the obtaining, verification or presentation of the contents” refers therefore to the resources and efforts that were called on to find, collect, verify and/or present existing materials. What constitutes a ‘substantial investment’ was explored in a case (<u>C-203/02 - The British Horseracing Board and Others</u>) in which the British Horseracing Board (and others) had objected to the re-use of the content of their database. Their case failed as the Court estimated that the collection and the presentations of the horseracing previsions and results did not require a substantial investment and, in consequence, the extraction and reuse of data was regarded as not being in contravention of database protection. This case is relevant to biology as many databases take pre-existing digital information from other sources and organize the data using widely accepted standard metadata, ontologies, and identifiers. Increasingly, biodiversity-oriented data environments (such as Catalogue of Life <u>Global Biodiversity Information Facility</u>, <u>Biodiversity Heritage Library</u>, <u>International Plant Name Index</u>, <u>Encyclopedia of Life</u>, or <u>Ocean Biogeographic Information Service</u>) rely to some extent on the content contributed by other databases or by individuals, projects and organizations. Such databases are likely to be ineligible for database protection and the use of some of the content of European biodiversity databases is likely to be legitimate. The value of such databases lies not in their content, but on the extent to which they are maintained to be current and accurate.

### 4.4 Protection against unfair competition

Many countries have legal regulations which seek to prevent unfair competition in industrial and commercial matters. The minimum standard for this protection, applicable in 194 countries, is defined by the Paris Convention for the Protection of Industrial Property, established in 1883 and amended most recently in 1979 (http://www.wipo.int/wipolex/en/details.jsp?id=12633). Art. 10^bis^ of the convention defines as prohibited unfair competition:

1. all acts of such a nature as to create confusion by any means whatever with the establishment, the goods, or the industrial or commercial activities, of a competitor;
2. false allegations in the course of trade of such a nature as to discredit the establishment, the goods, or the industrial or commercial activities, of a competitor;
3. indications or allegations the use of which in the course of trade is liable to mislead the public as to the nature, the manufacturing process, the characteristics, the suitability for their purpose, or the quantity, of the goods.”

Many countries consider that one form of unfair competition is to reproduce and exploit a competitor’s product or service which is ready for marketing without contributing any novel performance or investment. This legal protection does not aim at a defined intellectual property right, but at lawful commerce. It’s actions prevent behavior that could harm fair competition in an open market.

With respect to scientific images, it might constitute unfair competition to reproduce published images and sell them for individual profit. Unfair competition protection only applies if there is competition between the publisher of the images and the seller. The competition law does not prevent the utilization of published images for other non-competing purposes, such as for any scientific use.

### 4.5. Specific photograph protection in some European countries

A few European countries such as Germany and Austria have introduced special protection for photographs. The purpose is to protect against unfair competition. Photographers in these countries have a specific intellectual property right in their photographic production, but it applies only within that country. The right lasts for 50 years from the date of publication and protects against every form of re-use.

This special protection must be understood in the light of its historical background [84]. A revision of the Berne Convention was to take place in 1908 in Berlin. France asked for the extension of copyright protection to photographs. The German Reich was strictly opposed to this petition as it feared negative effects for its growing photographic industry. In order to prevent the French proposal, the German Reich introduced in 1907 this special protection for photographs granting the photographers fewer prerogatives than a copyright and lasting only 25 years. The Conference in 1908 ended with a compromise agreement that both solutions - copyright on one side, special protection on the other side - were acceptable in light of the Berne Convention. The result was that Germany did not protect photographs through copyright law. At the dawn of World War II, some countries under German influence (Austria, Denmark, Italy) followed their example. In 1948, the Berne Convention was revised again and at this time the copyright protection of photographs became compulsory. Instead of replacing the special protection with copyright protection, the aforementioned countries introduced the copyright protection for photographs into their national law, but also maintained the former protection. This double protection, referring to different kinds of photographs, was upheld also in later law revisions.

The specific photograph protection applies only to non-individual photographs, taken from three-dimensional objects. As it is the case in copyright law, the reprography of a print, a drawing or a pre-existing photography is not a photograph in the sense of these laws [85, N. 22 zu § 72 UrhG]. The protection is rather difficult to apply and has only little importance in practice. However, researchers working in one of these countries should be aware that the re-use of photographs under these legal systems is more problematic than in the rest of the world. Researchers not based in these countries, but wanting to use photographs from these countries, are not subject to this restriction.

## 5. Discussion

Considering this outline of intellectual property rights, we conclude that principles of copyright do not normally apply to scientific images because most images adhere to the conventions of the discipline. Certainly, copyright is not applicable to images that are intended to facilitate comparison among related taxa.

### 5.1 Rights in scientific images apply only in special cases

Copyrightable works are defined as individual, original human creations, that show originality in the form of presentation compared to other works of the same kind. Most scientific images lack an original form of presentation and so cannot qualify as copyrightable works. This is particularly true for machine-generated images, such as robotic systems used to digitize specimens in natural history collections, or pictures obtained from camera-traps positioned to monitor animal activity over time. Such pictures are not man-made and they can consequently not be copyrightable works. For the same reason, they do not qualify for the special photograph protection that applies within a few European countries.

Individually prepared photos and drawings, produced in line with scientifically recognized and standardized conventions, also fall outside the scope of copyright because of their standardized form of expression. Routine photographs and scans made from two-dimensional objects, as for example photos of herbarium sheets, are not copyrightable as they lack individuality and creativity (Fig. 14A).

Similar arguments apply to the combination of text and standardized images that make up taxonomic treatments. Treatments follow conventions to facilitate the effective documentation of facts, and comparison between descriptions. The expectations are so firm that peer review would not allow treatments that are individual in the sense of literature or art. They are technically “correct” if they are done according to the applicable protocols, and they are “incorrect” if they do not follow those standards. They express facts in a pre-established, standardized form. They do not, therefore, qualify as copyrightable works [86].

The same criterion leads to the conclusion that collections of biodiversity data are also not copyrightable by default as the selection and arrangements of those collections as well as their form of presentation follow predefined systems of biological classification, metadata, ontology, vocabulary and quantitative units. Tables of quantitative or qualitative information can be considered as collections of data, the selection and presentation of which are scientifically predefined. The more complete and systematic a collection, the less probable it is that it qualifies as a work in the sense of copyright. This conclusion does not devalue scientific work, but it is a logical consequence of copyright legislation that aims to protect individual forms of expression.

The situation is less consistent as far as wildlife illustrations are concerned. Some images are created for artistic purpose or to create a commercial product. Some photographs or drawings generated during field research and which are not produced in line with established standards, may fulfil the criterion of individuality and originality and therefore qualify as works in the sense of copyright. Copyright protection may apply to such images.

The situation may also be slightly different in E.U. countries which apply the *sui generis* protection for databases. Collections of biodiversity data may be subject to this specific protection against the re-use of a substantial part or the totality of the content of the database. Another exception that researchers should be aware of is the specific photograph protection in some European countries (such as Austria, Denmark, Germany, and Italy). Of course, these specific protection rules apply only in the countries that have introduced them. Outside these countries, the protection has no legal effect.

### 5.2 Blue2 - an updated “Blue List” -

‘The blue lists’ identify those elements which may reasonably be expected to occur in taxonomic works and, because of their compliance with conventions, lack the creativity that makes copyright applicable. The first list [12] addressed textual components in checklists, classifications, taxonomies, and monographs. Blue2 extends the list with 4 items relating to images. It is the view of the authors that the elements in the list below may be freely re-used unless restricted by a use agreement or a special limitation associated with a few countries. The original source of any re-used element should be cited, but this is demanded by the conventions of scholarship, not by legal obligation. The list may not be complete, and has not been tested in Court.

- A hierarchical organization (= classification), in which, as examples, species are nested in genera, genera in families, families in orders, and so on.
- Alphabetical, chronological, phylogenetic, palaeontological, geographical, ecological, host-based, or feature-based (e.g. life-form) ordering of taxa.
- Scientific names of genera or other uninomial taxa, species epithets of species names, binomial combinations as species names, or names of infraspecific taxa; with or without the author of the name and the date when it was first introduced. An analysis and/or reasoning as to the nomenclatural and taxonomic status of the name is a familiar component of a treatment.
- Information about the etymology of the name; statements as to the correct, alternate or erroneous spellings; reference or citation to the literature where the name was introduced or changed.
- Rank, composition and/or apomorphy of taxon.
- For species and subordinate taxa that have been placed in different genera, the author (with or without date) of the basionym of the name or the author (with or without date) of the combination or replacement name.
- Lists of synonyms and/or chresonyms or concepts, including analyses and/or reasoning as to the status or validity of each.
- Citations of publications that include taxonomic and nomenclatural acts, including typifications.
- Reference to the type species of a genus or to other type taxa.
- References to type material, including current or previous location of type material, collection name or abbreviation thereof, specimen codes, and status of type.
- Data about materials examined.
- References to image(s) or other media with information about the taxon.
- Information on overall distribution and ecology, perhaps with a map.
- Known uses, common names, and conservation status (including Red List status recommendation).
- Description and / or circumscription of the taxon (features or traits together with the applicable values), diagnostic characters of taxon, possibly with the means (such as a key) by which the taxon can be distinguished from relatives.
- General information including but not limited to: taxonomic history, morphology and anatomy, reproductive biology, ecology and habitat, biogeography, conservation status, systematic position and phylogenetic relationships of and within the taxon, and references to relevant literature.
- Photographs (or other image or series of images) by a person or persons using a recording device such as a scanner or camera, whether or not associated with light- or electron-microscopes, using X-rays, acoustics, tomography, electromagnetic resonance or other electromagnetic sources, of whole organisms, groups, colonies, life stages especially from dorsal, lateral, anterior, posterior, apical or other widely used perspectives and designed to show overall aspect of organism.
- Photographs (or other image or series of images) by a person or persons using a recording device such as a camera associated with light- or electron-microscopes, using X-rays, acoustics, tomography, electromagnetic resonance images or other electromagnetic sources) of parts of organisms such as but not limited to appendages, mouthparts, anatomical features, ultrastructural features, flowers, fruiting bodies, foliage, intra-organismic and inter-organismic connections, of compounds and analyses of compounds extracted from organisms that demonstrate the characteristics of an individual or taxon and/or allow comparison among individuals/taxa.
- Photographs (or other image or series of images) of whole organisms, groups, colonies, life stages, parts of organisms made by camera or scanner or comparable devices using automated procedures.
- Drawings of organisms or parts of organisms made by a person or persons to demonstrate the characteristics of an individual / taxon or to allow comparisons among taxa.
- Graphical / diagrammatic representation (such as, but not limited to, scatter plots with or without trend lines, histograms, or pie charts) of quantifiable features of one or more individuals or taxa for the purposes of showing the characteristics of or allowing comparison of individuals or taxa and made by a person or persons.

The first ‘Blue List’ [12] was based on the analysis of the prevailing law and usage patterns, involved a workshop, and input from the community. The analysis led to the conclusion that these elements were not copyrightable. We argue here that the same principle applies to drawings, photos, and maps that illustrate descriptions and circumscriptions of taxa, diagnostic characters, or any other element of the Blue2 list.

They do not qualify as copyrightable works as they are executed according to pre-established standards and protocols and are not individual in the sense of copyright.

The situation may differ as far as wildlife illustrations and photos produced during field research are concerned. Such illustrations may be expressed in an individual form and so qualify as works to which copyright may be applied.

### 5.3 Exceptions and limitations, fair use

Images that do not qualify as copyrightable work and that are not protected by any other intellectual property right, can be reused by any other person for any other legal purpose. Images and collections of images that are protected by copyright or by database protection may only be used with the consent of and under terms set by the rights holder. However, there are situations where even the use of copyrighted material is allowed without authorization.

The rules for these copyright exceptions vary fundamentally in different law systems. While the U.S. applies the so called “fair-use-defense”, European countries aim at the same objective through a catalogue of legally binding rules, called “exceptions and limitations”. In the U.K. and other common-law legislations, the exceptions and limitations are sometimes combined with a “fair-dealing-clause”. The different systems lead to different consequences with respect of the use of copyrighted material.

The “fair-use-clause” is part of the U.S. Copyright Act (17 U.S.C. § 107) and means that the unauthorized use of a copyrighted work will not be considered as copyright infringement if this use can be qualified as “fair”. In determining whether there is a fair use, the factors to be considered shall include the purpose and character of the use, including whether such use is of a commercial nature or is for nonprofit educational purposes, the nature of the copyrighted work, the amount and substantiality of the portion used in relation to the copyrighted work as a whole, and the effect of the use upon the potential market for or value of the copyrighted work. The function of the “fair-use-clause” is to give a case-by-case defense to persons who are sued for copyright infringement and where an objective consideration leads to the conclusion that such infringement was done in good faith or does not cause any relevant damage.

The “exceptions and limitations” which are used in the great majority of copyright laws are specific legal rules that authorize uses of copyrighted material for certain purposes. A commonly allowed exception to Copyright law is the use of copyrighted material for research purposes. These rules can be found in the national copyright laws and vary from country to country. The <u>E.U. Directive 2001/29/EC</u> tries to harmonize these rules for the E.U. Member States. It allows, amongst a whole catalogue of other elements, the Member States to provide in their national copyright law for exceptions and limitations for acts of reproduction made by publicly accessible libraries, museums or educational establishments as well as for acts of reproduction or communication for the purpose of illustration for teaching or scientific research. However, as has been illustrated by a recent investigation [79], despite this harmonization effort, national provisions in Europe on copyright and database protection regarding exceptions and limitations for research purposes differ not only in some details but also in substance.

The re-use of copyrighted material even for research purposes may therefore be hampered by current copyright and database protection. The risk is particularly true for international big data studies that were not foreseen by the law-makers and do not fit into the “fair-use”-criteria of U.S. copyright nor will be authorized by any exception rule of European copyright law. Such large projects are likely to inadvertently run counter to some exceptions and limitations or legislation that applies in some national jurisdictions.

### 5.4 No economic incentive

In creative fields, copyright is sometimes justified as a mechanism for encouraging innovation and creativity [87]. This is based on a very different model than that under which taxonomic researchers typically operate. Producers of creative content are economically incentivized directly by the appeal of their products and their marketability to consumers. Producers of scientific content, particularly in the context of articles for journals, are not incentivized in the same way. Researchers, often with support from public institutions and public or philanthropic grants, typically receive no direct financial incentive to create content. Recent experiments with financial incentivization for creators of scientific content have tended to increase the volume but not the quality of scientific content [88, 89]. To the contrary, journals often charge content creators a fee to defray costs associated with page layout, distribution, and other aspects of publishing. Until relatively recently, most journals also sought to acquire all intellectual property rights to the content that they published.

Because taxonomic research is funded in great part by public and philanthropic sources rather than capital investment, it follows that the fruits of this investment and labor are owed to the public rather than to investors. The current practice to cede legal rights to a private publisher, who may use these rights to restrict access to such publications, is highly problematic. The interests of both science and the public are better served by investing in publishing models that maximize accessibility, rather than producing research products paid for by, but not accessible to, the public [90]. Good science depends on independent scrutiny of reported results. When scientific reports survive scrutiny, their value increases. So, lowering access barriers to scientific content provides more opportunity for independent checks and leads to a healthier and more robust science, even when not publicly funded (e.g., citizen scientists). It is also in this context that legal principles concerned with the protection of creative content might not be properly applicable to scientific content.

## 6. Attribution

The principles of scholarship in taxonomic research include the expectation that relevant previous work be cited. Citation of publications identifies prior work and helps to assure reproducibility and comparability of the results of scientific research.

Citation offers a mechanism of providing credit to work by others, that is, attribution. In an increasingly digital world, we should be attentive to the principles of citation, comply with any legal obligations, and identify those who acquired the data or in any way contributed to the supply chain and/or added value to data [91].

### 6.1 Attribution in copyright

In the case of copyrighted work, citation is a legal obligation. As is stipulated in Art. 6^bis^ of the Berne Convention, every author shall have the right to claim authorship, “independently of the author’s economic rights, and even after the transfer of the said rights”. Nearly all states adhering to the Berne Convention have transformed this obligation into national law. This means that the name of authors must be joined to any use of the copyrighted work.

A special clause of the Berne Convention (Art. 10) deals with “quotations”. Quotations from a work made lawfully available to the public are permissible as long as the extent of the quotation does not exceed that justified by its purpose. Every quotation must be attributed to the source, and has to mention the name of the author if it appears in this source. This obligation is also transformed into the national law of nearly all member states of the Berne Convention and is therefore of general validity.

These legal obligations, however, apply only to copyrighted works or to quotations from copyrighted works. As we have seen before, scientific images are in most cases not copyrightable. As a consequence, there is no general obligation to attribute scientific images based in copyright law. Legal obligations are limited to the minority of cases where scientific images are copyrightable.

The E.U. database protection as well as the protection against unfair competition do not include any obligation to attribution. The same is true for the protection of non-copyrightable photographs as it exists in some European country. The only legal instrument that contains an obligation to attribute is therefore the copyright law.

Despite the absence of legal obligations, the tradition of citation has served science well, and benefits both the cited with credit and the citer with a reputation for integrity. It is the view of the authors that failing to recognize authors and/or suppliers of images is comparable to plagiarism. As noted by Patterson et al. [91], plagiarists have faced considerable sanctions such as having papers withdrawn, degrees retracted or dismissal from institutions.

### 6.2 How to attribute authorship in images

In the previous sections, we have laid out the arguments as to why images in scientific articles should be considered to be data, and not encumbered by copyright. We also argue that all previous work should be given attribution. Acceptance by the community that most images are not being subject to copyright must be accompanied attribution. It will be up to the scientific community to develop attribution standards.

In order to make recommendations about how to give attribution to the original authors, we take inspiration from a few other sources that have thought deeply about this subject, namely, the Digital Public Library of America (DPLA), Harvard University Library, and Europeana.

<u>The data use policy of the DPLA</u> is based on goodwill, not a legal contract. The DPLA is motivated by the belief that “the benefits of following (their) guidelines far exceeds any burdens and will foster the most creative and collaborative environment for the use/reuse of metadata from the DPLA.” As such, DPLA makes available all its metadata, also not subject to copyright for reasons similar to what we have argued in the preceding sections, under the <u>Creative Commons Zero (CC0) Public Domain Dedication.</u> CC0 permits use of the content for any purpose without having to give attribution. However, the DPLA wants to foster a community of practice that recognizes contributions, and giving attribution to all the sources of the metadata is crucial toward that objective. Thus, the DPLA recommends giving attribution to the data provider, all contributing data aggregators, as well as the DPLA itself. If, for any reason, attribution and rights information can’t be provided, DPLA suggests providing a link back to the relevant page on the DPLA website. Since data can change, DPLA recommends using the metadata via the DPLA API or via a hyperlink.

<u>Harvard University Library</u> provides bibliographic metadata under CC0 Public Domain Dedication. While Harvard does not impose any legally binding conditions on access to the metadata, they request that the user act in accordance with the following Community Norms of the Harvard Library with respect to the metadata. Specifically, Harvard requests that they, and OCLC and the Library of Congress, as appropriate, “be given attribution as a source of the Metadata, to the extent it is technologically feasible to do so.” They further request that any improvements made to the metadata be made available to everyone “without claiming any legal right in, or imposing any legally binding conditions on access to, the Metadata or your improvements, and with a request to act in accordance with these Community Norms.” The emphasis is not on legal obligations but on <u>community norms</u>.

Europeana, the digital portal for all of Europe’s culture, has a mission to “transform the world with culture!” As such, Europeana makes all metadata available on <u>europeana.eu</u> “free of restrictions, under the terms of the <u>Creative Commons CC0 1.0 Universal Public Domain Dedication</u>.” Europeana does encourage users to “follow the <u>Europeana Usage Guidelines for Metadata</u> and to provide attribution to the data sources whenever possible”.

Following in the footsteps of DPLA, Harvard Library, Europeana, and others, we recommend that authors recognize the author and source for each image that is used, and that publishers assign a DOI or other unique identifier to every image and mark the images with CC0. Publishers should provide a clear statement about copyright, recommend a suggested citation for images in the Terms of Use and the Data Policy sections of the website. Elsewhere we have argued that the use of unique identifiers with each data item (image in this case) allows the application of annotation technology that is capable of rewarding all members of the supply chain with credit and quantifiable metrics [91].

## 7. Acknowledgements

Thanks to Tim Smith (Zenodo, CERN), Lyubo Penev (Pensoft), Scott Miller (Smithsonian Institution), and Chuck Miller (Missouri Botanical Garden) for advice and discussion, and to the numerous colleagues we involved and challenged with this view over the last years in any conference, meeting, or gathering we attended.

